# Disentangling niche theory and beta diversity change

**DOI:** 10.1101/2021.07.05.451214

**Authors:** William Godsoe, Peter J. Bellingham, Elena Moltchanova

## Abstract

Beta diversity describes the differences in species composition among communities. Changes in beta diversity over time are thought to be due to selection based on species’ niche characteristics. For example, theory predicts that selection that favours habitat specialists will increase beta diversity. In practice, ecologists struggle to predict how beta diversity changes. To remedy this problem, we propose a novel solution that formally measures selection’s effects on beta diversity. Using the Price equation, we show how change in beta diversity over time can be partitioned into fundamental mechanisms including selection among species, variable selection among communities, drift, and immigration. A key finding of our approach is that a species’ short-term impact on beta diversity cannot be predicted using information on its long-term environmental requirements (i.e. its niche). We illustrate how our approach can be used to partition causes of diversity change in a montane tropical forest before and after an intense hurricane. Previous work in this system highlighted the resistance of habitat specialists and the recruitment of light-demanding species but was unable to quantify the importance of these effects on beta diversity. Using our approach, we show that changes in beta diversity were consistent with ecological drift. We use these results to highlight the opportunities presented by a synthesis of beta diversity and formal models of selection.

## Introduction

There is a need to understand the drivers of species diversity at broad spatial scales (Socolar et al. 2016, Jabot et al. 2020, Tatsumi et al. 2021). This is because many of the mechanisms that maintain species diversity operate at broad scales (Chesson 2000, Barabás et al. 2018, Usinowicz and Levine 2018), and anthropogenic activities reduce diversity at broad scales (Vellend et al. 2007, Baiser et al. 2012). Indices of diversity at broad spatial scales (gamma diversity) are divided into measures of average diversity within communities (alpha diversity) and differences among communities (beta diversity; see Table 1 for a list of terms). The more dissimilar communities are, the greater is beta diversity (Figure 1 A). Ecologists struggle to predict how beta diversity changes over time (Magurran and McGill 2011, McGill et al. 2015, Vellend 2016). For example, anthropogenic disturbance is hypothesized to homogenize communities, reducing beta diversity (Gutiérrez-Cánovas et al. 2013). However, anthropogenic disturbances (farming, selective logging, biological invasions, overhunting, and climate change) have been found to increase beta diversity in some studies and reduce it in others (Socolar et al. 2016); only urbanization has consistently reduced beta diversity. This suggests a need to better understand the mechanisms that drive beta diversity changes.

**Table 1.**
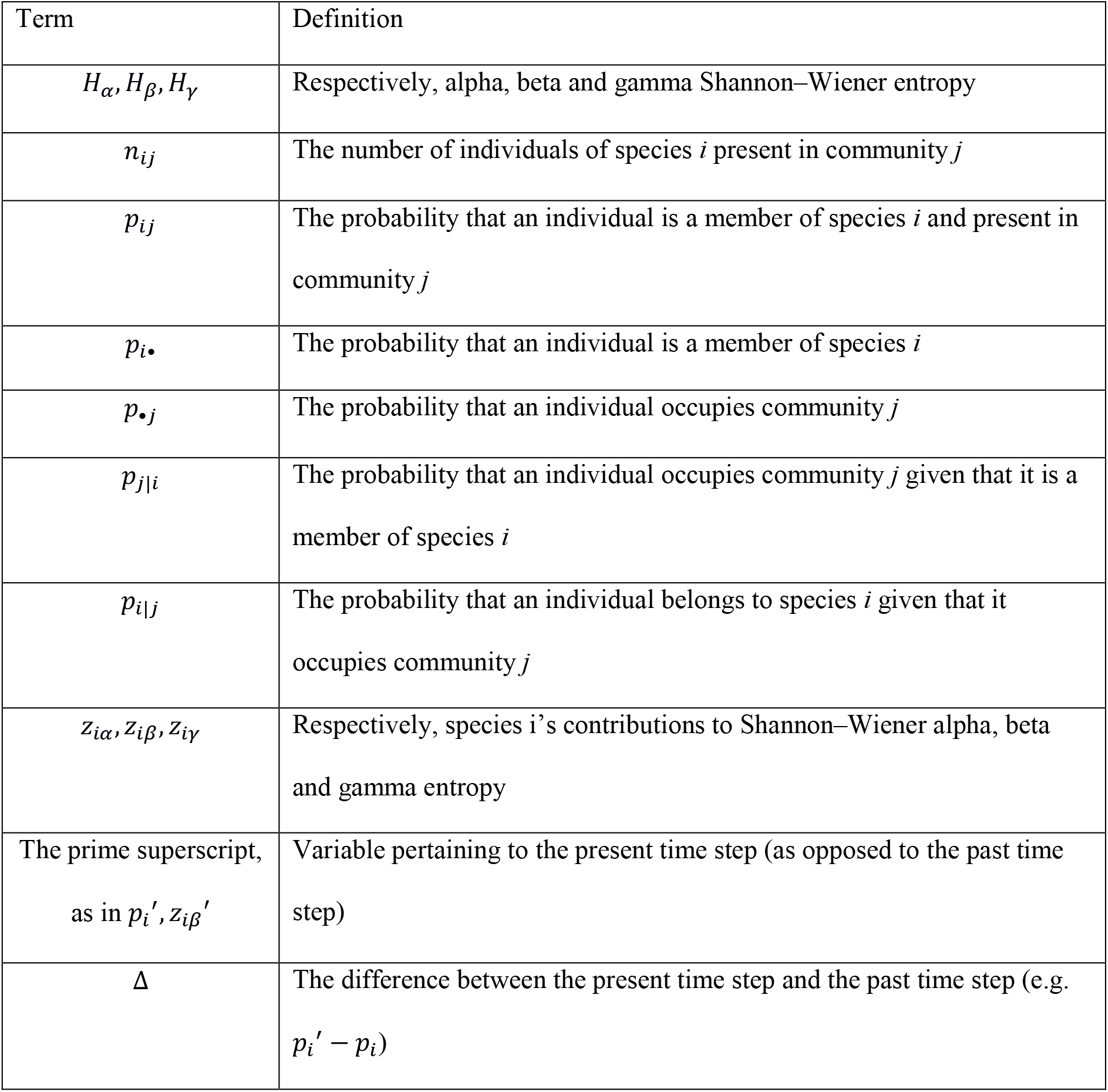
List of symbols used in the main text.

**Table 2.** List of terms used in the main text.

| Term | Definition |
| --- | --- |
| Alpha diversity | Diversity within communities, averaged across a metacommunity. |
| Average local rarity | A measure of how rare individuals of a given species are within their local community, averaged over all individuals. |
| Average metacommunity rarity | A measure of how rare individuals of a given species are in the metacommunity. |
| Average relative rarity | A comparison of the rarity of individuals of a given species in their local community, relative to the rarity of that species in the metacommunity. |
| Beta diversity | Diversity among communities. |
| Community | The set of individuals found within a single location. |
| Diversity index | A statistic that is proportional diversity (i.e. to some measure of the rarity of species). Unlike true diversity, different diversity indices typically have different units. |
| Drift | Change in a measurement of interest due to stochastic variation in the number of descendants produced by different types of organisms. |
| Gamma diversity | The diversity of species across a metacommunity. |
| Generalist | For our purposes, a species that is capable of establishing or persisting in many communities. |
| Metacommunity | The collection of communities being studied. |
| Niche | The set of communities that are suitable to a species at equilibrium; i.e. the set of communities where environmental conditions are suitable enough to allow the species to establish or persist (Holt 2009). |
| Specialist | For our purposes, a species that is unable to establish or persist in many communities. |
| True diversities | Measures of diversity with common units of “effective numbers of elements” (Jost 2007), calculated using Hill numbers (Hill 1973). |

**Figure 1.**
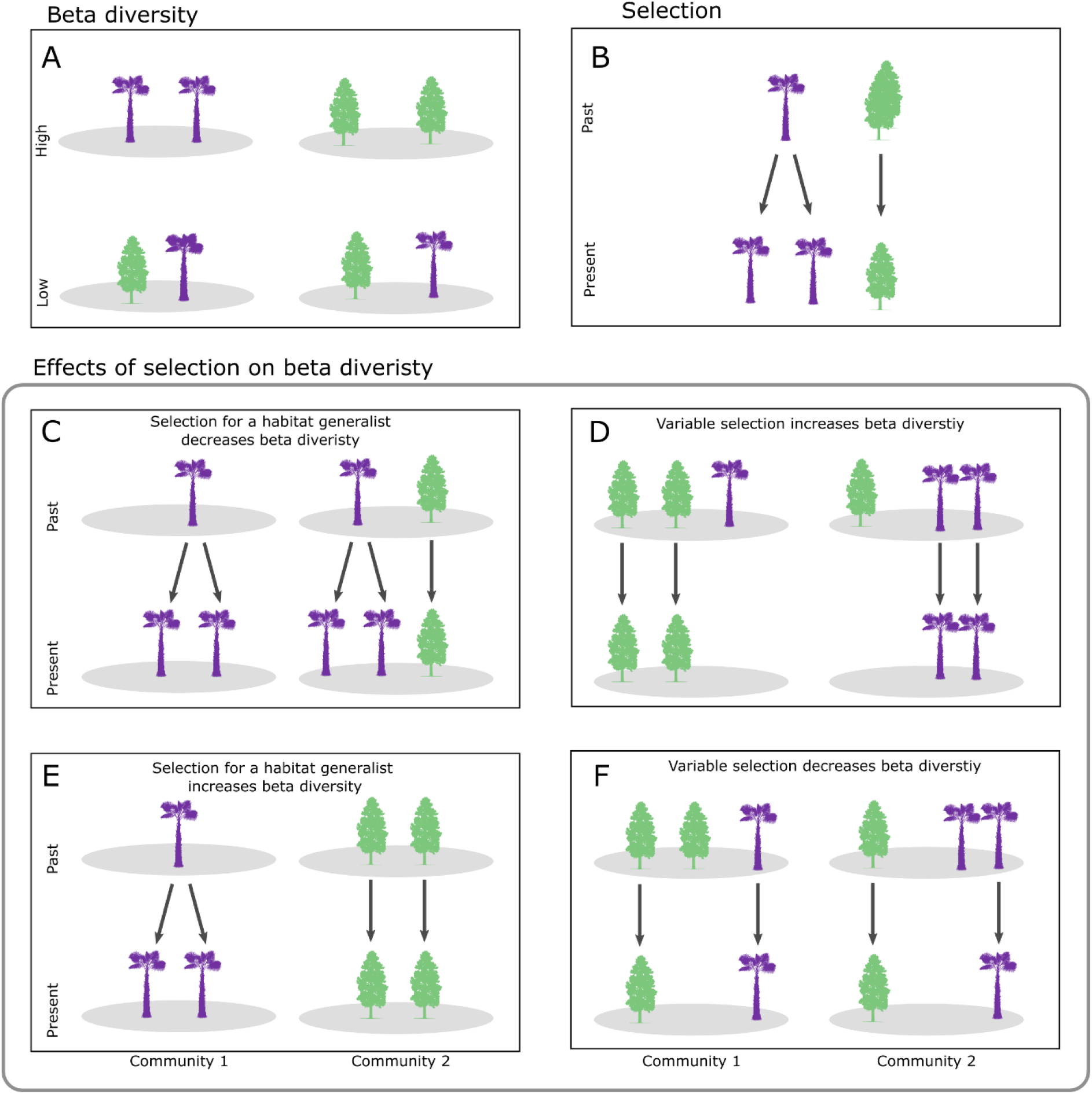
A) Beta diversity describes dissimilarities in ecological communities across space. It is highest when each community (grey ellipse) has a distinct species: a palm (purple) or a pine (green). Beta diversity is and lowest when the relative abundance of all species is the same in each community. B) Selection describes change over time due to the tendency for some organisms in the past (i.e. ancestors) to contribute more descendants than others. In this case, palm trees in the past produced more descendants than the pine trees. C) Illustrates the common hypothesis that selection in favour of a habitat generalist (the palm) at the expense of a habitat specialist (the pine) decreases beta diversity (here and in subsequent panels change in weighted Shannon–Wiener entropy: Δ*H*_*β*_ ≈−0.055). In this case, palms are initially distributed evenly across both communities and produce more offspring than the pine, leading to a community that is more homogeneous in the present than it was in the past. D) This panel illustrates the common hypothesis that variable selection increases beta diversity. In this case, selection favours the pine in community 1 and the palm in community 2 (Δ*H*_*β*_ ≈0.636). However, selection sometimes has counterintuitive effects on beta diversity. E) This illustrates a case where selection in favour of a habitat generalist increases beta diversity. Here the palm species is a habitat generalist that can potentially thrive in both communities, but dispersal limitation has limited it to community 1. The palm produces more descendants, and these descendants remain in the same community as their parents, so beta diversity increases (Δ*H*_*β*_ ≈0.057). F) This illustrates a case where variable selection decreases beta diversity (Δ*H*_*β*_ ≈−0.056): the palm is favoured in community 1 and the pine in community 2.

Beta diversity is thought to change due to selection based on species’ niches (i.e. its environmental requirements at equilibrium; Holt 2009). Selection based on species’ niche characteristics is thought to lead to predictable changes in beta diversity (Chase 2003, Chase and Leibold 2003, Hawkins et al. 2015, Myers et al. 2015, Tucker et al. 2016, Vellend 2016). This idea is based on the insight that fitness differences among species are analogous to fitness differences among genotypes in evolutionary biology (Day 2005, Vellend 2010, Mallet 2012). Selection, in this context, would result when one species increases in frequency by producing more descendants than others (Figure 1B). Selection that favours habitat generalists (those that can establish or persist in many communities) can reduce beta diversity (Figure 1C), such as when urbanization favours the same non-native species in many locations (Chase 2007, Lepori and Malmqvist 2009, Hawkins et al. 2015, Myers et al. 2015, Catano et al. 2017, Thorn et al. 2020).

Niche differences can lead to selection in favour of a species in some communities and against it in others (hereafter spatially variable selection), and this is expected to increase beta diversity (Figure 1D; Vellend 2016, Stubbington et al. 2019). Ideally we would use information on what Hutchinson (1957) called a species’ realized niche, the set of environments that are suitable to a species in the absence of dispersal among environments (Godsoe et al. 2017). Though the term realized niche can have other connotations (Hutchinson 1978, Soberón and Peterson 2005). However, niche-based prediction of beta diversity rarely measure environmental suitability directly, instead, they typically rely on observations of species’ current distributions. A species’ current distribution is often a poor surrogate for a species’ realized niche (MacArthur 1972, Pulliam 2000). For instance, dispersal limitation can restrict a species to a subset of its realized niche. This is the case for the grass *Vulpia fasciculata* which occurs along beaches in southern Great Britain. Transplant experiments show that locations well to the north of its current range are a part of its niche (Norton et al. 2005).

Conversely, current distribution may overestimate a realized niche when dispersal is easy, such that species occur in environments unsuitable for them (e.g., several Himalayan plant species disperse to and colonize high-elevation communities in which they cannot persist; Klimeš and Doležal 2010). Current distributions may also overestimate the realized niche in the case of long-lived species persisting in communities outside their current niche (e.g., low-elevation populations of beech *Fagus sylvatica* in Spain that likely colonized when the climate was cooler; Peñuelas et al. 2007, Jump et al. 2009). Such mismatches between niches and distributions are common, occurring in 54% of transplant experiments (Hargreaves et al. 2014).

Mismatches between species’ niches and distributions can lead to misleading predictions of changes in beta diversity over time. For example, many introduced species are generalists, which are ultimately capable of thriving across different communities. However, many species’ introductions start with a few propagules reaching a few communities in a new region, so that they experience a “lag phase”, where they are rare and restricted to a subset of suitable communities (Davis 2009, Aikio et al. 2010). During this lag phase, the success of generalists can increase beta diversity. For example, Figure 1E shows a species with broad environmental tolerances, meaning that it can persist in both communities. Although it is a habitat generalist, previous dispersal limitation has restricted it to a single community. In that community its population grows. A second species is a specialist in community 2 and, although it cannot persist in community 1, it still occurs there extremely rarely due to dispersal. Over time, as the population of the habitat generalist grows, beta diversity increases (See Appendix S 1 for example calculations). This contrasts with the widespread theoretical prediction that the success of habitat generalists reduces beta diversity (Chase 2007, Lepori and Malmqvist 2009, Hawkins et al. 2015, Myers et al. 2015, Catano et al. 2017, Thorn et al. 2020).

Niche theory also predicts that spatially variable selection can increase beta diversity (Vellend 2016, Stubbington et al. 2019). Spatially variable selection operates when one species is favoured in one community and disfavoured in another (figure 1 D). In practice, spatially variable selection can reduce beta diversity when it favours species in communities where they are rare. This can happen when dispersal barriers between communities break down. When the biogeographic barrier breaks down, small numbers of each species can invade communities occupied by the other species (Figure 1F). This leads to spatially variable selection with newly arrived species succeeding in communities where they are rare. Though examples as clear cut as figure 1F may be rare in nature, this form of spatially variable selection is supported by phylogenetic analyses of terrestrial vertebrates (Pigot and Tobias 2013, Pigot and Tobias 2015), and can reduce beta diversity.

We propose a novel solution to resolve discrepancies between predictions based on niche theory and observed changes in beta diversity over time through a formal measurement of selection. In evolution, the standard way to measure selection is to specify the attribute we are studying, then determine if that attribute is correlated with fitness (Price 1970, Lande and Arnold 1983, Price 1995, Frank 2012a, Queller 2017). For example, to learn whether selection among mammals increases average body size, we measure species’ body sizes and determine if larger mammals have greater fitness than smaller mammals (Rankin et al. 2015). Selection among species in a metacommunity can be measured similarly. To know if selection increases diversity, we should measure species’ contributions to diversity, then determine if these contributions predict a species’ relative fitness. In a previous study we showed that change in many diversity indices results from selection based on species’ rarity within a single community (Godsoe et al. In press). To measure selection’s effects on beta diversity we would need to quantify each species’ contribution to beta diversity. Because beta diversity is typically interpreted as a property of communities (Jost 2007, Ellison 2010), not species, we develop a revised approach inspired by tools from information theory (Cover and Thomas 2012).

We illustrate how selection’s effects can be measured using the beta diversity index associated with Shannon–Wiener entropy (*H*_*β*_), an index commonly used in ecology (Magurran 2013), with deep mathematical links to information theory and analyses of selection (Shannon 1948, MacArthur 1965, Cover and Thomas 2012, Frank 2012b). A particular species’ contribution, *H*_*β*_ is its average relative rarity, a measure of the difference between its rarity in the metacommunity and its rarity in individual communities. Average relative rarity is high for species that are rare in some communities and common in others. We show that change in beta diversity can be partitioned into fundamental mechanisms using the Price equation (Price 1970). These observations are then generalized to multiplicative partitions of true diversities expressed in common units (Jost 2006, 2007). Traditional diversity indices, such as richness and Shannon–Wiener entropy, are difficult to compare directly because each index use different units (MacArthur 1972, Hill 1973). As a result, it is now common practice to convert these indices into “true diversities” with common units– the number of equally common species requited to produce the observed diversity index for alpha and gamma diversity and the number of equally diverse distinct communities for beta diversity (Jost 2006, 2007, Sherwin et al. 2017, Gaggiotti et al. 2018). Throughout, we highlight how analyses of selection leads to different predictions from niche theory and how these predictions could be tested.

## The model

### Species’ contributions to beta diversity

We consider a metacommunity consisting of communities *j* = 1, …, *N* occupied by species *i* = 1, …, *S*, where the number of individuals of species *i* in community *j* is *n*_*ij*_ (see Table 1 for a list of terms and definitions). The total number of individuals across the metacommunity is ∑_*i*_ ∑_*j*_ *n*_*ij*_ and the probability that an individual belongs to species *i* and resides in community *j* 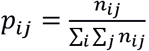. The choice of what counts as a single individual will of course alter measurements of diversity, but our methods work regardless of this choice. Similarly, it could be convenient to define *n*_*ij*_ using other units such as biomass, but this choice does not change our derivations. To measure selection we must keep track of the relative abundance of each species in the metacommunity, which is defined as 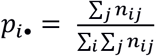, and the relative abundance of individuals belonging to species *i* given we are in community *j*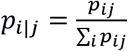. To weight the contributions of multiple communities it is useful to define the probability that an individual belongs to community *j*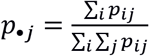, and the probability that an individual is in community *j* given that it belongs to species *i*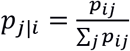.

Shannon–Wiener entropy is a natural starting point for analyses of selection, which are often defined as a partition of changes in average of measurements (Queller 2017, Lion 2018). This average is commonly defined on an additive scale (Rice 2004, Lion 2018). Shannon–Wiener entropies are measured on the same additive scale that is used to measure selection (Jost 2006, Frank 2012b). For Shannon–Wiener entropy, rarity is defined as the negative log of species’ relative abundance (Jost 2006). An individual’s contribution to gamma diversity is its species’ rarity across the metacommunity: −*log*(*p*_*i*•_). An individual’s contribution to alpha diversity is its rarity score within its local community: −*log*(*p*_*i*|*j*_). An individual’s contribution to beta diversity is its relative rarity (Figure 2); i.e. the difference in its rarity score in the metacommunity versus its rarity score in its local community: 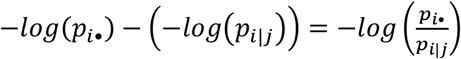. A species’ contribution to *H*_*β*_ is the average relative rarity of all individuals of species *i* in the metacommunity:

**Figure 2.**
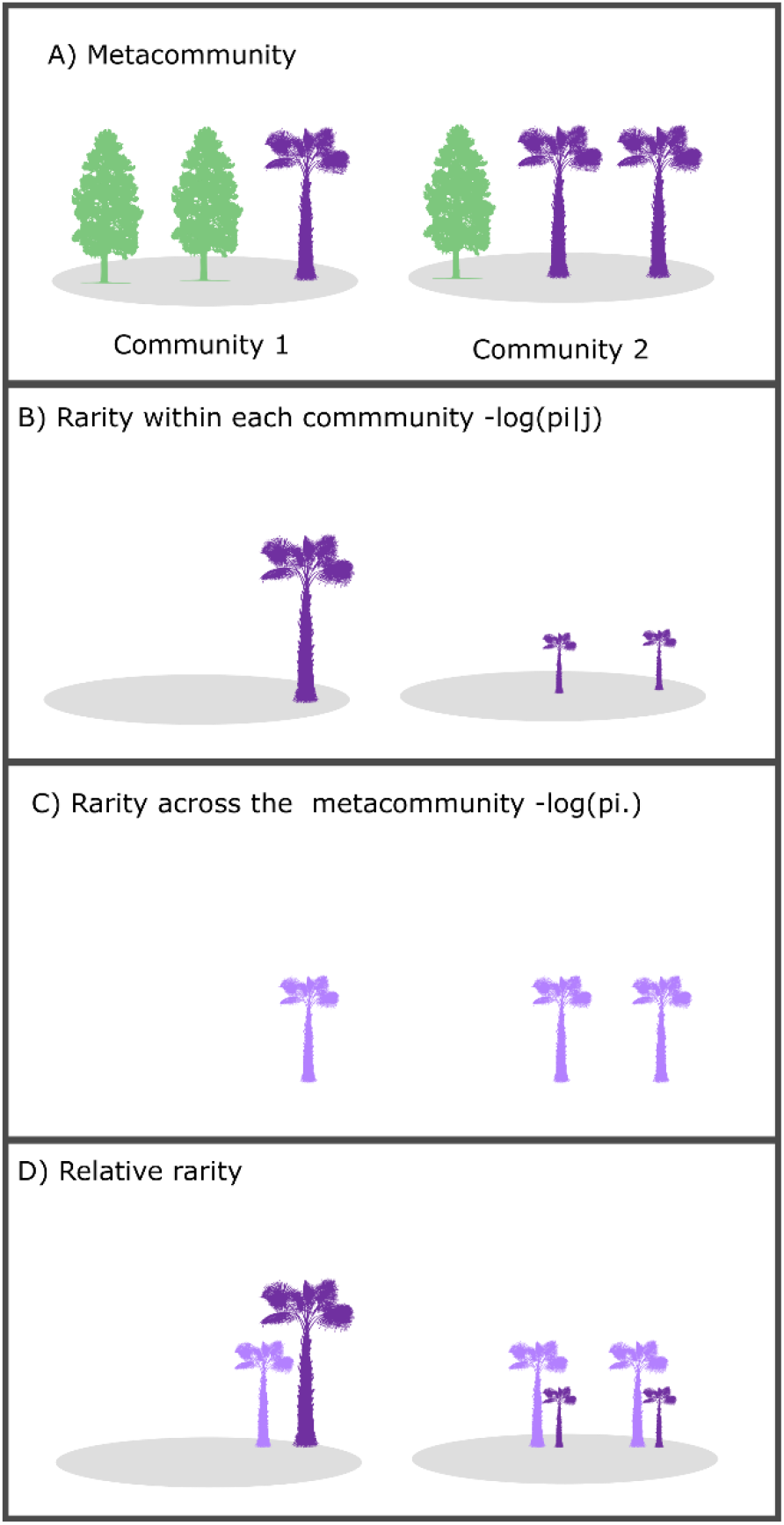
A) An example of a metacommunity consisting of two species, a palm (purple) and a pine (green) distributed across two communities. B) Focusing on one species (the palm), we can measure rarity within each community. To emphasize this, trees here and in subsequent panels have been scaled in proportion to rarity scores. In community 1 the palm is the rarer species (*p*_*i*|*j*_ = 1/3, leading to a local rarity score of –log(1/3)=1.10). In community 2 the palm is less rare (*p*_*i*|*j*_ = 2/3, leading to a local rarity score of –log(2/3)=0.41). The palm’s contribution to alpha diversity depends on average local rarity: *z*_*iα*_ = ∑_*j*_ *p*_*j*|*i*_ (−*log*(*p*_*i*|*j*_)) = 1/3×1.10 + 2/3 × 0.41 = 0.64. C) The metacommunity rarity score for individual palms is −*log*(*p*_*i*•_) = −*log*(1/2)=0.69. The palm’s contribution to gamma diversity is an average of metacommunity rarity scores *z*_*iγ*_ = ∑_*j*_ *p*_*j*|*i*_(0.69)=0.69. D) The per-individual contribution to beta diversity is the difference between local rarity scores and metacommunity rarity scores. We can visualize this as the difference in height when the palm is scored according to its local rarity (dark purple) versus when it is scored according to its metacommunity rarity (light purple), *z*_*iβ*_ = ∑_*j*_ *p*_*j*|*i*_ ((−*log*(*p*_*i*•_)) − (−*log*(*p*_*i*|*j*_))) = 1/3 × (0.69– 1.10) + 2/3 × (0.69–0.41) = 0.056).

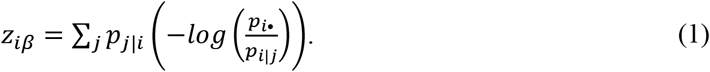

Average relative rarity reaches a minimum of 0 when a species has the same relative abundance in all communities. *z*_*iβ*_ can be arbitrarily large when a species is exceptionally rare in some communities but common in others. Measurements of selection typically weight the contributions of all individuals equally. Consistent with this, Equation 1 assumes that all individuals are weighted equally, regardless of the number of individuals in their local community. Appendix S2.1 equation 16 provides an alternative expression for alternative weightings.

In turn, *H*_*β*_ is an average of each species’ contribution, weighted by its relative abundance in the metacommunity.

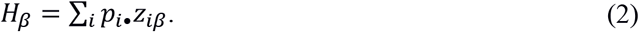

Each species’ contribution to *H*_*γ*_ and *H*_*α*_ can be defined using an equation analogous to Equation 2 as *z*_*iα*_ and *z*_*iγ*_ respectively (see Appendix S 2.1).

### Species’ contributions to changes in beta diversity

We define change in *H*_*β*_ over time as the difference between *H*_*β*_ in the present and *H*_*β*_ from a time in the past:

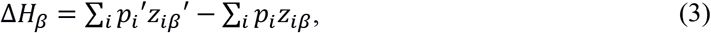

where the present time period is indicated with primes (i.e. 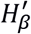) and Δ indicates the difference between one time period and another. This definition can be applied over any timescale (i.e. it may be 1 hour or 1000 years earlier). We will refer to individuals in the past as ancestors and individuals in the present as descendants (Frank 2012a). Equation 2 is agnostic about what came before the past time step, meaning that diversity in the past time step may be due to a combination of processes, including selection, drift, speciation and immigration (Vellend 2016). We assume that individuals may disperse from one community to another, but, for now, we assume that the metacommunity as a whole is closed to immigration. Equation 3 can be partitioned into two fundamental mechanisms; the proof is provided in the derivation for Equation 1 in Frank (2012a).

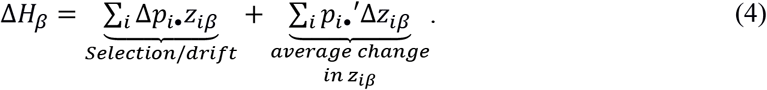

This is the simplest way to express one of the most fundamental theorems in evolution: the Price equation (Frank 2012a, Queller 2017, Frank and Godsoe 2020). We can obtain analogous partitions for alpha or gamma diversity by substituting *z*_*iβ*_ with species’ contributions to alpha or gamma diversity.

The “selection/drift” term measures change in *H*_*β*_ resulting from change in species’ relative abundance between the present and past 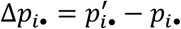. The selection/drift term is positive when individuals belonging to species with high *z*_*iβ*_ scores leave the most descendants, and negative when individuals belonging to species with low *z*_*iβ*_ scores leave the most descendants. This term can be interpreted as the covariance between relative fitness and *z*_*iβ*_ (Frank 2012). In evolutionary theory the selection/drift term is commonly called “selection”, because in many mathematical models the number of descendants left by a given species depends only on its relative fitness. In nature this term can be interpreted as a measure of the combined effects of selection and drift among species (Rice 2004). In a subsequent section we suggest a null model to tease apart drift’s contribution.

The selection term in Equation 4 treats all descendants equally, regardless of the community they occupy. By implication, immigration within the metacommunity does not change Equation 4, because descendants make the same contribution regardless of the community they occupy. (See Appendix S 2.4 for illustrations of this principle).

The change in one species’ contribution to beta diversity is the difference between *z*_*iβ*_ in the present and past 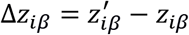. The “average change in *z*_*iβ*_” term measures the effect of these changes across all species. This term is positive when individual species become more patchily distributed over time. This can be due to geographically variable selection; for example, when habitat specialists decrease in relative abundance in unsuitable habitat communities and increase in suitable communities. Changes in *z*_*iβ*_ can also be due to dispersal (Appendix S 2.4). For example, passive dispersal may move species from communities where they are common to communities where they are rare, evening out their distribution and, over time, reducing *z*_*iβ*_.

Simplified examples help to illustrate how our framework distinguishes selection among species from other mechanisms. Consider, for example, the metacommunity in Figure 1D, where beta diversity has increased over time. Across the metacommunity there is no selection among species because the relative abundance of each species remains unchanged. This can be verified with Equation 4, where the relative abundance of palms (as in Figure 1D) remains 1/2 (Δ*p*_*palm*•_= 0) and the relative abundance of pines remains 1/2 (Δ*p*_*pine*•_= 0), leading to a selection term of 0 (i.e. ∑_*i*_ Δ*p*_*i*•_*z*_*iβ*_ = ∑_*i*_ 0 × *z*_*iβ*_ = 0). In other words, the fitness of each species is equivalent when averaged across the metacommunity. The change in *H*_*β*_ is average change in *z*_*iβ*_ scores, weighted by species’ relative abundances. Each species has succeeded in one community and failed in the other.

Calculations for past *z*_*iβ*_ are provided in (Figure 2; *z*_*palmβ*_ = *z*_*pineβ*_ = 0.056). Present scores for palms are given by 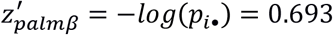, which is found by re-arranging Equation 2 to: 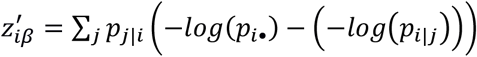 then substituting in 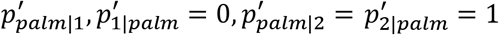 and 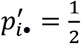. Note that, by convention, 0log0 is set to 0 (Cover and Thomas 2012). A similar calculation reveals that 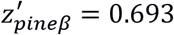. From this we can compute change in *z*_*iβ*_ for each species: 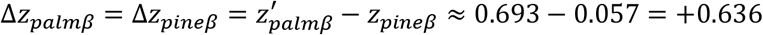. The average change in *z*_*iβ*_ is 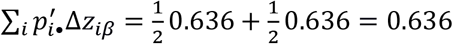. The effect of selection + change in *z*_*iβ*_ = 0+ 0.636. This is equal to the total change in *Hβ* (Appendix S1).

This partition of change in *H*_*β*_ can be converted into an analysis of change in true Shannon– Wiener diversity (Jost 2006, 2007). Shannon–Wiener diversity is measured on a multiplicative scale, and so it is useful to define a new measure of change over time on a multiplicative scale, the ratio of present diversity 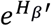 to past diversity 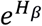. Using the quotient rule for exponents (as in Jost 2006 equation 5a,b), this can be re-written as:

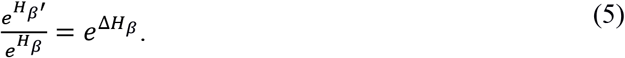

We can then substitute in Equation 4, giving:

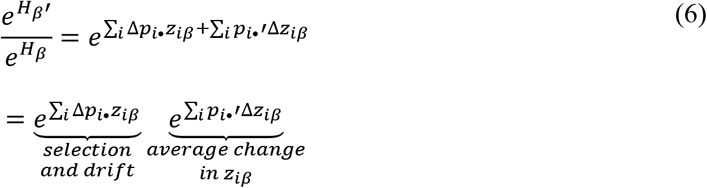

That is, the ratio of Shannon–Wiener beta diversity in the present, relative to the past, is decomposed into the multiplicative effects of selection and average change in *z*_*iβ*_. Equation 6 can be illustrated with the example in Figure 1 D. When explaining Equation 4 we measured the effect of selection and drift on *H*_*β*_: 0, and the effect of changes in *z*_*iβ*_ on *H*_*β*_: 0.636. Substituting these values into Equation 6 gives e^0^e^0.636^≈1.89. This is equal to the ratio of Shannon–Wiener diversity in the present to diversity in the past: 1.89 (Appendix S1). All true beta diversities (Jost 2006, 2007, Tuomisto 2010) can be partitioned in a manner similar to Equation 4, but at the cost of more mathematical abstraction (Frank and Godsoe 2020; Appendix S 2.2).

Equations 4 and 6 assume that the metacommunity is closed immigration, but an extension of the Price equation can be used to relax this assumption (Kerr and Godfrey-Smith 2009, Frank 2012a). This approach distinguishes two categories of individuals in the present community: 1) individuals that are descended from members of the past community, 2) immigrants (i.e. individuals that are not descended from past members of the community). Much as in Equation 4, descendants’ contributions to *H*_*β*_ change are partitioned into selection and average change in *z*_*iβ*_. The effect of these mechanisms is weighted by the relative abundance of descendants in the present community, meaning that these mechanisms matter less when immigration is common. The effects of immigration depend on the proportion of individuals in the present community that are immigrants, and the difference between the *z*_*iβ*_ scores of immigrants and the *z*_*iβ*_ scores of ancestors (Appendix S 2.3).

### Partitioning beta diversity change in nature

We illustrate how our approach can be used to measure the causes of diversity change in nature using data from a montane tropical forest during recovery from hurricane damage (Appendix S3). Tanner (1977) sampled forests in four sites within 25–175 m of each other that differed in topography and soil nutrient availability on and near the main ridge of the Blue Mountains of Jamaica (18°05′ N, 76°39′ W; 1580–1600 m). The Mor ridge site (hereafter Mor) has low soil nutrient availability, with low alpha diversity and several habitat specialists (Tanner 1977). The three other sites are more fertile, include many habitat generalists, and have higher alpha diversity: Col forest, Wet Slope forest, and Mull Ridge forest (hereafter Col, Slope and Mull, respectively; Tanner and Bellingham 2006). At each site all tree individuals ≥3 cm in diameter at 1.3 m height (dbh) were identified, tagged, and measured during an initial survey in 1974.

These sites were re-censused in 1984, then hit by Hurricane Gilbert in 1988―considered the most powerful hurricane in the Caribbean in the 20th century―and then re-censused in 1994 (Tanner and Bellingham 2006). The hurricane caused increased tree mortality, both immediately and over several years as a result of stem damage (Bellingham et al. 1992, Tanner et al. 2014), and widespread defoliation, increasing the light available in the forest floor for approximately three years (Bellingham et al. 1996). As a result, alpha diversity increased in some sites, which Tanner and Bellingham (2006) hypothesized was due to the recruitment of light-demanding species, but they did not quantify the contributions of these to overall species diversity.

Using counts of individual trees in each of the Jamaican forest sites, we computed *H*_*β*_ change between the 1984 sampling period before the 1988 hurricane and the 1994 sampling period after it. Tanner and Bellingham (2006) found some species in the 1994 census that had not been recorded in 1984 and represented recruitment. These individuals were < 3 cm in diameter before 1994. Few would have geminated since the 1988 hurricane because diameter growth rates were low (Tanner et al. 2014). The success of these individuals represent recruitment, but in the Price equation they are treated as immigrants since none of their ancestors were in the community in 1988 (Kerr and Godfrey-Smith 2009).

We treated all other individuals as descendants. This choice probably leads to some bias because a small fraction of the individuals we treat as descendants were probably immigrants. However, we expect this bias to be negligible. To alter the partitioning of diversity substantially, immigrants must be are common and the distribution of individuals that are immigrants must be dissimilar to the distribution of ancestors (Equation 28 in Appendix S 2.3.). Experiments suggest that immigrants to the Mor site are uncommon because species outside this site are excluded by the nutrient-poor soil (Sugden et al. 1985). Immigrants may be more common in the other three sites, but the distribution of immigrants in these three sites is probably similar to the distribution of ancestors, since all three sites are within broadly similar forests (Tanner 1977).

The observed changes in diversity between 1984 and 1994 are just one possible realization of stochastic changes in abundance across the metacommunity. To assess the uncertainty associated with this stochasticity we used parametric bootstrapping. To do this we assumed that the 1984 counts observed for each species in each site are fixed, then simulated counts for species *i* in community *j* in 1994 using realizations from a Poisson probability distribution with a mean equal to the observed present count of species *i* in location *j*. A total of 10,000 simulated datasets was obtained. From these we estimated empirical 95% confidence intervals for total diversity change in alpha, beta, and gamma diversity, along with change due to selection/drift, average change in *z*_*i*_ (species’ contribution to diversity), and immigration. One limitation of this procedure is that it estimates our confidence in the combined effects of selection and drift, rather than distinguishing their individual contribution.

To distinguish drift’s effects from selection, we developed a null model where all individuals present in 1984 were equally likely to produce descendants censused in 1994. This model used a Monte Carlo simulation based on the actual datasets. To do this, the 1984 count of species *i* in community *j* was fixed as observed. The 1994 counts of species *i* at location *j* were simulated using a Poisson distribution with a mean equal to 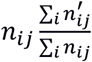 (where 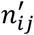 is the present count of species i in community j). We then used 10,000 replicates to calculate a 95% empirical confidence interval of change expected due to drift: ∑_*i*_(*p*_*i*•|*D*_ − *p*_*i*_)*z*_*iβ*_, which can be compared to the analogous portion of the selection/drift term in Equation 28 in Appendix S 2.3.

## Results

In the 1984 pre-hurricane survey, *H*_*β*_ across the four Jamaican forests was 0.53. Individuals of some species, such as *Cyrilla racemiflora*, are approximately equally rare across communities, leading to a low *z*_*iβ*_ score (Figure 3A). Species that were unusually rare in more fertile sites had higher *z*_*iβ*_ scores, such as *Lyonia octandra* (Figure 3B), a habitat specialist (Tanner 1977). Some of the highest *z*_*iβ*_ scores are from rare species, such as *Cinnamomum montanum* (Figure 3 B), which is represented by only two individuals in the Col site. Given the small number of individuals observed, the distribution of *C. montanum* individuals may simply represent chance rather than niche specialization (Thorn et al. 2020).

**Figure 3.**
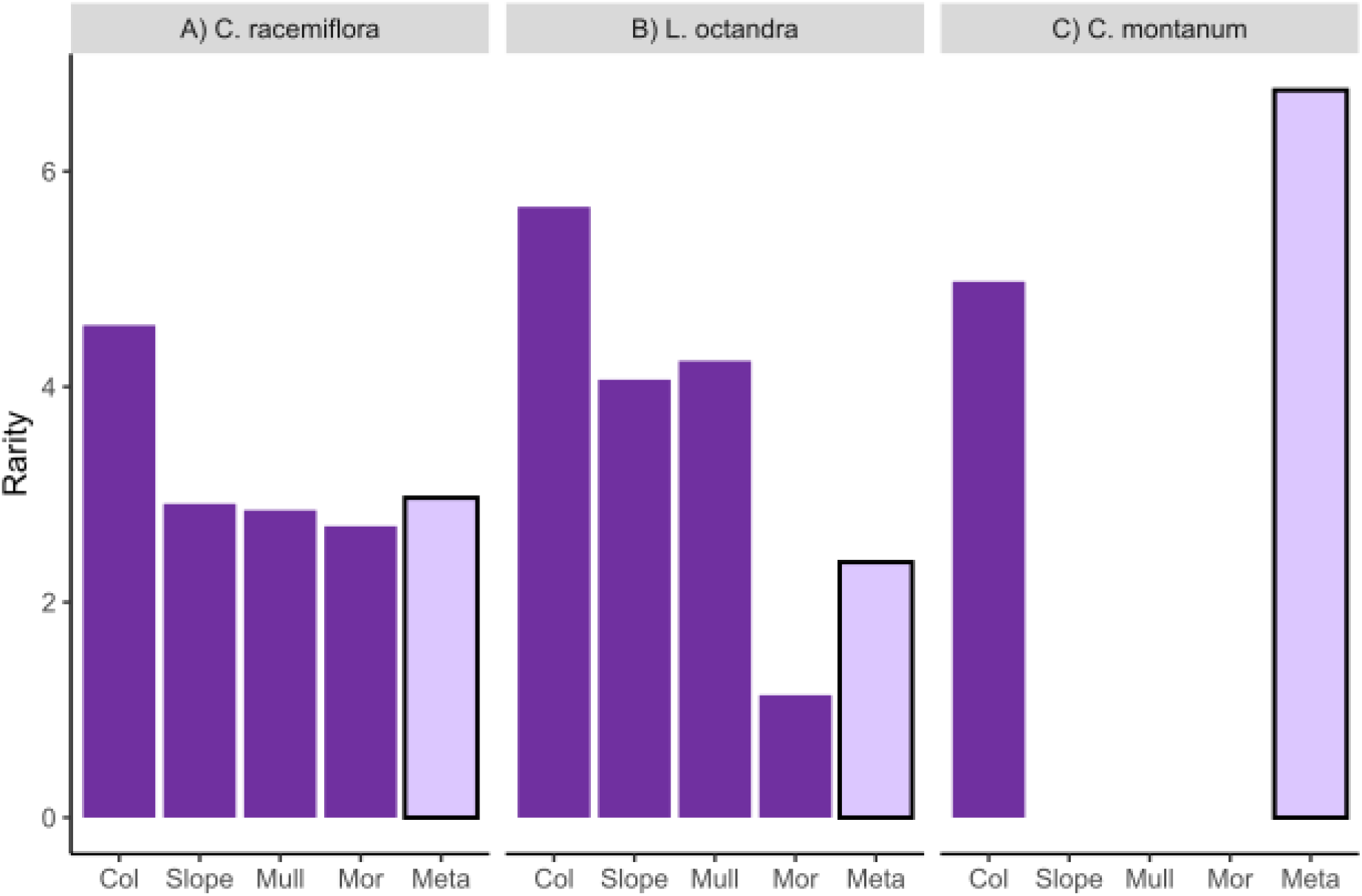
Comparisons of local rarity scores −*log*(*p*_*i*|*j*_) (dark purple) and metacommunity rarity scores (−*log*(*p*_*i*•_) (light purple) for three tree species across sites in Jamaican montane rain forests. A) *Cyrilla racemiflora* is a habitat generalist with similar rarity scores in each of the communities and across the metacommunity. This species has a low *z*_*iβ*_ score (*z*_*iβ*_ = 0.094). B) *Lyonia octandra* is considered a habitat specialist on nutrient-poor soils. This species is rare in the three nutrient-richer sites and more common in the nutrient-poor site (Mor). *Lyonia octandra* has a high *z*_*iβ*_ score (*z*_*iβ*_ = 0.921). C) Of the three species, *Cinamonum montanum* has the highest species *z*_*iβ*_score (*z*_*iβ*_ = 1.77). There were only two individuals of this species, both in the Col site. Since only two individuals were observed, their distribution probably represents chance rather than niche requirements. See Figure 2 for explicit calculations of *z*_*iβ*_.

*H*_*β*_ was essentially unchanged between 1984 and 1994 (ΔH_β_ = -0.0006). Recruitment of new species increased *H*_*β*_, although the effect was weak (Figure 4), in part because the new recruits represent only ∼1.3 % of the present community. Species with high *z*_*iβ*_ scores were roughly as likely to increase in relative abundance as species with low *z*_*iβ*_ scores, leading to a small combined effect of selection and drift (Figure 4). The change in beta diversity among the sites was likely to be a result of drift alone, because we did not detect an effect of selection with a significance level set to 0.05 (p = 0.38, null model expectation: -0.016 to 0.017).

**Figure 4:**
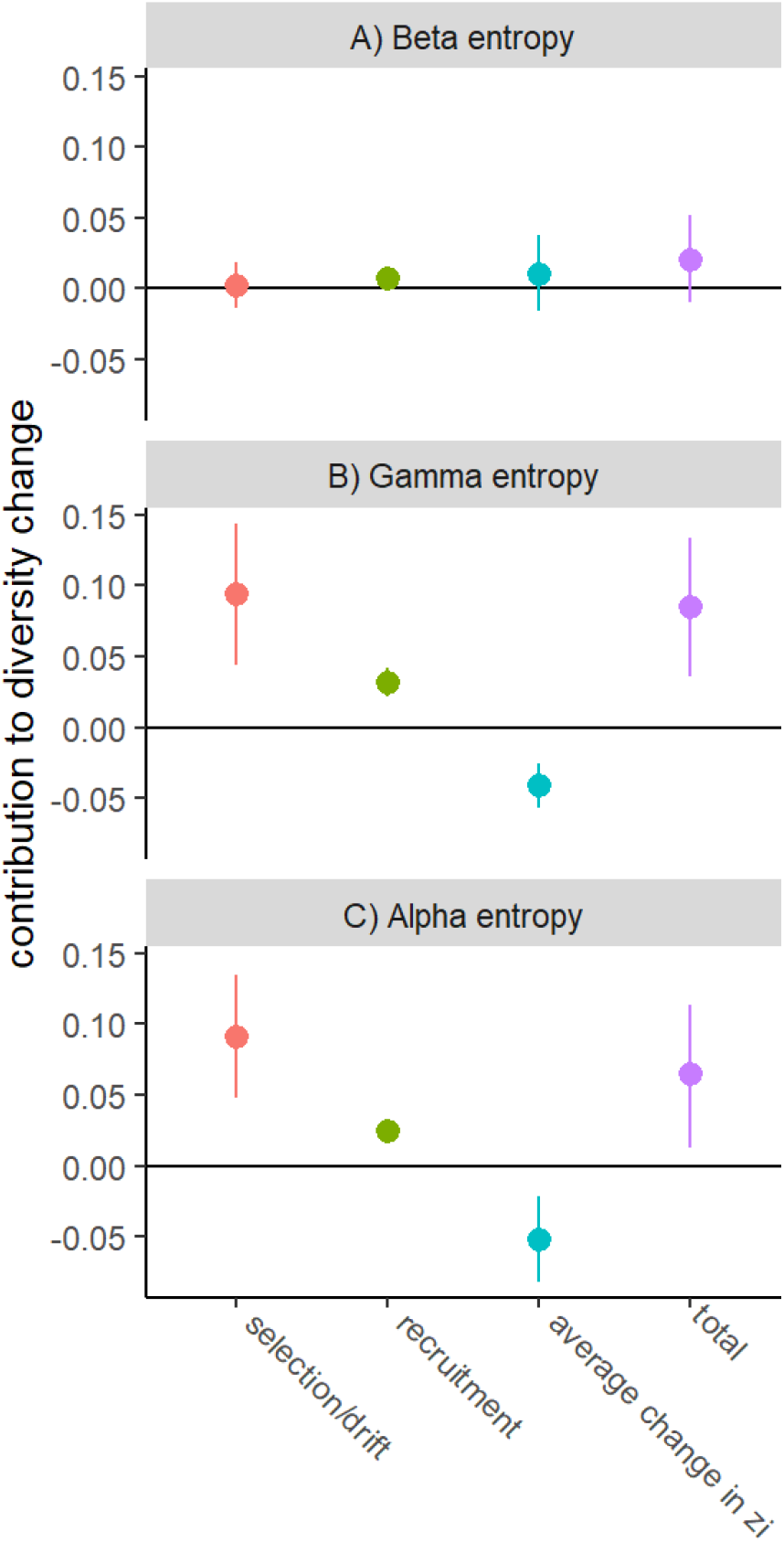
Partitions of the causes of for the four sites in Jamaican montane rain forests between 1984 and 1994 in A) Shannon–Wiener beta entropy, B) gamma entropy, and C) alpha entropy. Error bars represent boot-strapped 95% confidence intervals.

*H*_*γ*_ increased substantially between 1984 and 1994 (ΔH_γ_ = 0.104), influenced most strongly by selection and drift (Figure 4). The observed association between z_iγ_ and the relative abundance of descendants is stronger than would be expected by chance alone, indicating that selection contributed to this increase in *H*_*γ*_ (p < 0.05). The selection term represents an additive contribution of individual species with *Urbananthus critoniformis* making the largest contribution. This is likely to be light-demanding (Bellingham, personal observation), but was not previously identified as a key component of change in diversity (Tanner & Bellingham 2006). Recruitment of new species also increased *H*_*γ*_. Change in *H*_*α*_ was qualitatively similar to change in *H*_*γ*_ (Figure 4).

## Discussion

This work is a synthesis of the theory of selection and beta diversity change. The synthesis relies on finding each species’ contribution to beta diversity and measuring the association between these contributions and relative fitness. We show how these tools can lead to better predictions than niche theory (Figure 1), and how they can be used to measure beta diversity change in nature (Figure 4). Using these tools we show that beta diversity change in Jamaican montane rainforests is consistent with drift while change in alpha and gamma diversity is consistent with selection. Both selection and beta diversity represent rich topics, and they cannot be merged entirely in a single paper. Instead, we have focused on the most tangible insights that lie at the intersection of these two topics.

Our framework simplifies some of the ambiguities that are caused by dispersal. For example, the effect of dispersal within the metacommunity is captured entirely by the change in *z*_*iβ*_, not selection. The effect of dispersal from outside the metacommunity can be captured using modified versions of the Price equation (Appendix S 2.3). This separation is easiest when dispersal can be measured directly, such as when immigrants belong to new species (Tatsumi et al. 2021), in experiments where dispersal is manipulated (Cadotte and Fukami 2005), or in marine systems where ocean circulation simulations are used to model larval dispersal (Bode et al. 2011, Watson et al. 2011). When unmeasured dispersal is likely, its effects could be estimated using parametric bootstrapping. To do this, individual simulations would assign members of the present community to be immigrants in proportion equal to the expected probability of immigrants from outside the metacommunity. The selection, change in *z*_*iβ*_, and immigration terms can then be recalculated, including these sources of uncertainty. Based on the natural history of montane forests, we expect our analyses of diversity change in Jamaican rainforests to be robust to this bias (Tanner 1977, Tanner 1982, Sugden et al. 1985).

Our approach focuses on individual’s contributions to beta diversity instead of the equilibrium dynamics emphasized by niche theory (Hutchinson 1957, MacArthur 1972, Chase and Leibold 2003, Holt 2009). Equilibrium dynamics often provide a misleading picture of the distribution of individuals (Pulliam 2000, Hargreaves et al. 2014). The limitations of niche theory become clearer when we recognize that beta diversity change can be partitioned into a component that depends on change in species’ relative abundance, and a component that depends on change in the distribution of individuals across communities (Equation 4). These sources of change are difficult to predict using niche theory alone. Short-term changes in relative abundance need not reflect equilibrium dynamics, and hence species’ niches (Hastings 1981, 2004, Van Cleve and Feldman 2008).

It may help to highlight cases where niche theory is most likely to provide a misleading guide to beta diversity change. Niche theory predicts that selection that favours habitat generalists will decrease beta diversity (Myers et al. 2015, Vellend 2016). Our partitioning predicts that selection in favour of widespread species will decrease beta diversity. Niche theory will be wrong when specialists are widespread. This can happen when habitat destruction causes time-delayed extinctions (i.e. extinction debt; Tilman et al. 1994, Kuussaari et al. 2009). Niche theory will also be wrong when generalists are narrowly distributed. This can happen when a small population of a generalist species establishes in a new region, such as during an invasion (Wilson and Lee 1989). Niche theory predicts that spatial variability in selection increases beta diversity (Myers et al. 2015, Vellend 2016). Our partitioning predicts that variable selection across communities can decrease beta diversity when species are favoured when rare (Figure 1F). There are, of course, situations where equilibria are a useful guide to diversity change. In these cases, our method may still be useful as it provides tools to quantify the contributions of individual mechanisms and species to diversity change.

Our re-analysis of Tanner and Bellingham (2006) shows that the change in *H*_*β*_ is no more than we would expect by drift alone. This implies that species with high *z*_*iβ*_ scores were no more likely to leave descendants than were species with low *z*_*iβ*_ scores. This is surprising because Tanner and Bellingham (2006) suggest that one group of species with high *z*_*iβ*_ scores were “most resistant to hurricaine damage”; the habitat specialists in the nutrient-poor Mor site. Beta diversity in other ecosystems disturbed by hurricanes is also mainly influenced by drift (Chen et al. 2020). Tanner and Bellingham (2006) emphasized the recruitment of new species in the post-hurricane survey, but were unable to quantify its effects. We were able to show that this had a statistically significant, but biologically small, effect on beta diversity (Figure 4).

Although change in beta diversity was due primarily to drift, selection had a substantial effect on alpha and gamma diversity. This indicates that species which made a large contribution to alpha diversity had an advantage, species that made a large contribution to gamma diversity had an advantage but species which made a large contribution to beta diversity had no advantage. This lack of selection on beta diversity is surprising. It runs contrary to previous predictions that that selection, operating in a similar way across local communities (i.e. selection on alpha and gamma diversity), will to homogenize community composition (Catano et al. 2017).

Given that our goal is primarily to illustrate our approach and its interpretation, we elected to present results from time intervals that bracket Hurricane Gilbert (1984–1994). This time interval is discussed extensively by Tanner and Bellingham (2006), who reported other sampling periods but note dramatic changes between 1984 and 1994, including defoliation and immediate mortality. In a preliminary analysis we found that change in *H*_*γ*_ was larger in the sampling interval of 1984 and 1994 than in the other sampling intervals 1974–1984 and 1994–2004. In other applications there will certainly be advantages to studying *H*_*β*_ change over longer time periods. The 10-year sampling interval we use is short relative to the lifespan of the trees studied, meaning that selection probably reflects which trees survived the hurricane rather than reproduction post-hurricane.

There will also be opportunities to improve statistical inference in our framework. The bootstrapping and Monte Carlo procedures we used assume large and representative sampling. This assumption is reasonable for our re-analysis of Tanner and Bellingham (2006), a study that identified all stems in their plots. In other datasets these assumptions will be limiting. Many measures of diversity, including Shannon–Wiener entropy, are biased when sampling is incomplete (Chao and Shen 2003, Marcon et al. 2014). When this is the case it will be desirable to incorporate one of the bias-corrected estimators available (Chao et al. 2013, Chao and Jost 2015).

There is a long-standing debate over whether beta diversity can be used to study community assembly (Anderson et al. 2011). Our work suggests that beta diversity is linked to community assembly, but these links can be missed by traditional analyses. Predictions of beta diversity change often focus on the contributions made by each species’ niche, but this is only an imperfect surrogate for species’ contributions to beta diversity. Recognizing these contributions leads to new tools to predict and measure beta diversity change.

## Supporting information

example code folder

## Acknowledgements

We thank Edmund Tanner for establishing the permanently marked sites in Jamaica, for data collection, and for helpful comments on earlier drafts. Thanks are due to these authorities in Jamaica: (i) the Heads of the Department of Forestry in the Ministry of Agriculture, and the Park Managers of the Blue and John Crow Mountains National Park for permission to undertake the study; (ii) the Directors of the Botanic Gardens Division of the Ministry of Agriculture for permission to use the Cinchona Botanic Gardens as a field station; (iii) G. H. Sidrak, L. B. Coke, P. V. Devi Prasad, R. Robinson, D. Webber. E. Hyslop, M. Webber, J. Cohen and K. McLaren of the Department of Botany (more recently Life Sciences) of the University of the West Indies for assistance and facilities. Hannah Scrase and John Healey provided valuable field assistance. PJB was funded by the Strategic Science Investment Fund of the New Zealand Ministry of Business, Innovation and Employment’s Science and Innovation Group. Invaluable comments were provided by two anonymous reviewers, Zach Marion, Sarah Flanagan, Philip E. Hulme, Sarah Wyse, Dominique Gravel, Ray Prebble and Jennifer Bufford.

## Notes

### Competing Interest Statement

The authors have declared no competing interest.

### Summary of Updates

This version incorporates a few minor suggested changes such as additional citations on how we define a species' niche and the casues of some of the scenarios we describe.

