## Supplemental appendix 1 for "Disentangling niche theory and beta diversity change"

# Revising for resubmission on 26/02/2021

#this version tries to follow the calculations in the order presented int the text.

#other than tidying code i am changing the way that zibeta is calculated, this does not change the numerical value
#but the interpretation of the current calculation may be clearer for ecologists.


rm(list=ls())
#require(vegan)
require(vegetarian)

#This function modifies the log function in R so that our calculations follow
#the convention in information theory that 0*log(0)=0,
#by default 0*log(0) returns NaN, 
logMod<-function(X){
  Y<-log(X) #currently set to base e
  Y[X==0]<-0
  Y[is.infinite(Y)]<-0
  return(Y)
}


calculateDeltaD<-function(past, pres, divOrder, equalW=F){
  
  #this is the calculation when communities are weighted equally.
  if(equalW==T){
    return(d(pres,lev="beta",q=divOrder)-d(past,lev="beta",q=divOrder))
  }
  
  
  pjpast<-rowSums(past)/sum(past)
  pjprest<-rowSums(pres)/sum(pres)
  
  d(pres,lev="beta",wts=pjprest,q=divOrder)-d(past,lev="beta",wts=pjpast,q=divOrder)
}

calculateDeltaH<-function(past,pres){
  pjpast<-rowSums(past)/sum(past)
  pjprest<-rowSums(pres)/sum(pres)
  
  #d returns jost's "true" beta diversity (i.e. e^(Hb))
  #this result must be logged to obtain Hb

  log(d(pres,lev="beta",wts=pjprest))-log(d(past,lev="beta",wts=pjpast))
}


#this is an interpretation from an older draft where zibeta is interpreted as species metacommunity divergence. 
zibetaFOld<-function( meta=NA){
  
  pj<-rowSums(meta)/sum(meta)
  pi<-colSums(meta)/sum(meta)
  pjGiveni<-t(t(meta)/colSums(meta))
  piGivenj<-meta/rowSums(meta)
  
  
  colSums(pjGiveni*logMod(pjGiveni/pj))
}


#this computes zibeta using a revised interpretation of zibeta as average relative rarity
zibetaF<-function( meta){
  
  pj<-rowSums(meta)/sum(meta)
  pi<-colSums(meta)/sum(meta)
  pjGiveni<-t(t(meta)/colSums(meta))
  piGivenj<-meta/rowSums(meta)
  
  
  #this suggests that the sign convention has switched accidentally which will need to be checked.
  colSums(pjGiveni*(t(-t(-logMod(piGivenj))+(-logMod(pi)))))
}


##Compact code to compute selection and transmission terms
divPart<-function(past, pres1){
  
  #summary statistics needed to compute hbeta
  pjPast<-rowSums(past)/sum(past)
  piPast<-colSums(past)/sum(past)
  pjGiveniPast<-t(t(past)/colSums(past))
  pj<-rowSums(pres1)/sum(pres1)
  pi<-colSums(pres1)/sum(pres1)
  pjGiveni<-t(t(pres1)/colSums(pres1))
  
  
  #zibeta scores using equation 2 from the main text
  ziBetaPast<-colSums(pjGiveniPast*logMod(pjGiveniPast/pjPast))
  ziBeta<-colSums(pjGiveni*logMod(pjGiveni/pj))
  
  
  Selection<-(pi-piPast)%*%ziBetaPast
  Transmission<-pi%*%(ziBeta-ziBetaPast)
  
  return(c(Selection=Selection, Transmission=Transmission, Total=Selection+Transmission))
}


############Figure 1 delta Hb for each panel##########################################


#this is panel c in the current figure seleciton for a generalist decreases beta diversity
past<-cbind(s1=c(1,1), #abundance of species 1 in each patch
            s2=c(0,1)) #abundance of species 2 in each patch


pres<-cbind(s1=c(2,2), #abundance of species 1 in each patch
            s2=c(0,1)) #abundance of species 2 in each patch


calculateDeltaH(past,pres) 

#For details on how shannon wiener entropy is computed see this function which calculateDeltaH
#re-organizes raw scores to  weighted shannon wiener diversity using the d() function in
#the vegetarian package, then converts back to entropy


#this is panel d in the current figure variable selection increases beta diversity
past<-cbind(s1=c(1,2), #abundance of species 1 in each patch
            s2=c(2,1)) #abundance of species 2 in each patch


pres<-cbind(s1=c(0,2), #abundance of species 1 in each patch
            s2=c(2,0)) #abundance of species 2 in each patch


calculateDeltaH(past,pres)


#this is panel e in the current figure selection in favor of a habitat generalist increases beta diversity
#using the version of shannon wiener entropy weighted by the number of individuals per community used throughout the rest of the text


past<-cbind(s1=c(1,0), #abundance of species 1 in each patch
            s2=c(0,2)) #abundance of species 2 in each patch


pres<-cbind(s1=c(2,0), #abundance of species 1 in each patch
            s2=c(0,2)) #abundance of species 2 in each patch


calculateDeltaH(past,pres)


#this is panel F in the current figure: variable selection decreases beta diversity

past<-cbind(s1=c(1,2), #abundance of species 1 in each patch
            s2=c(2,1)) #abundance of species 2 in each patch


pres<-cbind(s1=c(1,1), #abundance of species 1 in each patch
            s2=c(1,1)) #abundance of species 2 in each patch


calculateDeltaH(past,pres)


###########################Effect of weighting on figure 1E and figure 1F##############################################

#############Introduction show that "several beta diversity metrics increase for Figure 1 E" ###########################


past<-cbind(s1=c(1,0), #abundance of species 1 in each patch
            s2=c(0,2)) #abundance of species 2 in each patch


pres<-cbind(s1=c(2,0), #abundance of species 1 in each patch
            s2=c(0,2)) #abundance of species 2 in each patch


#here are metrics that increase when individuals are weighted equally (i.e. when communities are weighted unequally)
calculateDeltaD(past,pres,0, equalW=F) #richness
calculateDeltaD(past,pres,1, equalW=F) #shannon
calculateDeltaD(past,pres,2, equalW=F) #simpsons
calculateDeltaD(past,pres,3, equalW=F) #hill number of order 3, i.e. one that weights common species more heavily than simpsons


#note, a common recommendation is to  weight communities equally (and weight individuals unequally). When this is done, for the example presented above
#diversity does not change. 

calculateDeltaD(past,pres,1, equalW=T)

#This is an artifact of making communities where abundances are small whole numbers (i.e. 0, 1,2).
#To see this modify the scenario slightly, Assume that dispersal leads to the presence of a small 
#number of species 2 (the specialist) in community 1.
#This preserves the qualitative pattern (i.e. species 1 is a generalist and is favored), species 2 is a specialist
#but now beta diversity metrics increase when communities are weighted evenly.


pastM<-cbind(s1=c(1,0), #abundance of species 1 in each patch
            s2=c(0.01,2)) #abundance of species 2 in each patch


presM<-cbind(s1=c(2,0), #abundance of species 1 in each patch
            s2=c(0.01,2))

#now, as in the weighted case the success of a generalist increases diversity
calculateDeltaD(pastM,presM,0, equalW=T) #richness
calculateDeltaD(pastM,presM,1, equalW=T) #shannon
calculateDeltaD(pastM,presM,2, equalW=T) #simpsons
calculateDeltaD(pastM,presM,3, equalW=T) 


############Introduction show that "beta diversity is intermediate in the past and as small as possible in figure 1 F"##########


past<-cbind(s1=c(1,2), #abundance of species 1 in each patch
            s2=c(2,1)) #abundance of species 2 in each patch


pres<-cbind(s1=c(1,1), #abundance of species 1 in each patch
            s2=c(1,1)) #abundance of species 2 in each patch


#here are metrics that increase when individuals are weighted equally (i.e. when communities are weighted unequally)
calculateDeltaD(past,pres,0, equalW=F) #richness
calculateDeltaD(past,pres,1, equalW=F) #shannon
calculateDeltaD(past,pres,2, equalW=F) #simpsons
calculateDeltaD(past,pres,3, equalW=F) #hill number of order 3, i.e. one that weights common species more heavily than simpsons


#For this example the results are unchanged when communities are weighted equally and individuals are weigthed unequally.
calculateDeltaD(past,pres,0, equalW=T) #richness
calculateDeltaD(past,pres,1, equalW=T) #shannon
calculateDeltaD(past,pres,2, equalW=T) #simpsons
calculateDeltaD(past,pres,3, equalW=T) #hill number of order 3, i.e. one that weights common species more heavily than simpsons


###########Figure 2 example calculations for zigamma alpha and beta referred to in model #################################


past<-cbind(s1=c(1,2), #abundance of species 1 in each patch
            s2=c(2,1)) #abundance of species 2 in each patch


#summary statistics needed to compute hbeta
#pj<-rowSums(past)/sum(past)
pi<-colSums(past)/sum(past)
pjGiveni<-t(t(past)/colSums(past))
piGivenj<-past/rowSums(past)


#the rarity of a species in a given community is:
-logMod(piGivenj)

#the rarity of a species across the meta community is:
-logMod(pi)


#this is the calculation for ziAlpha associated with panel B
ziAlpha=colSums(pjGiveni*(-logMod(piGivenj)))

#zigamma is just -logMod(pi), but to ease comparisons between zialpha, zigamma and zibeta it is useful to note
#that zigamma can be intepreted as a weighted average as well

ziGamma=colSums(pjGiveni*(-logMod(pi)))


#Zibeta is the difference between these rarity scores averaged. This calculation is a bit awkward in r so see zibetaF function for formula
zibetaF(past)
 
#Another way to interpret zibeta is as the difference between zigamma and zialpha
ziGamma-ziAlpha


###########Full calculations for Figure 1 D ##############################################


#this is panel d in the current figure variable selection increases beta diversity
past<-cbind(s1=c(1,2), #abundance of species 1 in each patch
            s2=c(2,1)) #abundance of species 2 in each patch

#summary statistics needed to compute hbeta
pj<-rowSums(past)/sum(past)
pi<-colSums(past)/sum(past)
pjGiveni<-t(t(past)/colSums(past))
piGivenj<-past/rowSums(past)
zibeta<-zibetaF(past)

pres<-cbind(s1=c(0,2), #abundance of species 1 in each patch
            s2=c(2,0)) #abundance of species 2 in each patch


#summary statistics needed to compute hbeta
pjP<-rowSums(pres)/sum(pres)
piP<-colSums(pres)/sum(pres)
pjGiveniP<-t(t(pres)/colSums(pres))
piGivenjP<-pres/rowSums(pres)
zibetaP<-zibetaF(pres) #this calculation is also explained in the main text


#Confirm that frequencies do not change across the metacommunity

piP-pi


#confirm that there is no selection across the metacommunity

selection=(piP-pi)%*%zibeta

#Change in zibeta scores (for each per species)
zibetaP-zibeta

#change in zibeta (averaged over the metacommunity)
transmission=(zibetaP-zibeta)%*%piP

#Total change
selection+transmission

#is equal to change in shannon wiener entropy

calculateDeltaH(past, pres)


###########Convert Figure 1 D from shannon wiener entropy to shannon wiener diversity#################################


#the ratio of present shannon wiener diversity to past shannon wiener diversity is equal to
d(pres, lev="beta", wts=pjP)/d(past, lev="beta", wts=pj)

#the multiplicative effects of exp(selection) and exp(transmission)
exp(selection)*exp(transmission)


###################################to delete###################################


####################make a simplified figure################################


past<-cbind(s1=c(1,2), #abundance of species 1 in each patch
            s2=c(1,1)) #abundance of species 2 in each patch


#summary statistics needed to compute hbeta
pj<-rowSums(past)/sum(past)
pi<-colSums(past)/sum(past)
pjGiveni<-t(t(past)/colSums(past))
piGivenj<-past/rowSums(past)


#the rarity of a species in a given community is:
-logMod(piGivenj)

#the rarity of a species across the meta community is:
-logMod(pi)


#Zibeta is the difference between these rarity scores averaged. This calculation is a bit awkward in r so see zibetaF function for formula

ziBeta<- zibetaF(past)


#Hbeta is
pi%*%ziBeta


#Check that this is the same as the standard calculation of Hbeta using the vegetarian package

log(d(past,lev="beta",wts=pj))


#d calculates the equivalent number of evenly distributed communitiees needed to produce
#the observed beta diveristy score. To find shannon wiener beta diversity, this must be log-transformed


########################second figure illustrating change in beta diversity############################
Hpast<-log(d(past,lev="beta",wts=pj))
piPast<-pi
ziBetaPast<-ziBeta


pres1<-cbind(s1=c(0,3), #abundance of species 1 in each patch
            s2=c(2,0)) #abundance of species 2 in each patch


#summary statistics needed to compute hbeta
pj<-rowSums(pres1)/sum(pres1)
pi<-colSums(pres1)/sum(pres1)
pjGiveni<-t(t(pres1)/colSums(pres1))


#the information associated with each species in each patch is:
-logMod(pjGiveni)

#the information associated with all individuals in each patch is:
-logMod(pj)


#compute contributions to beta diversity
ziBeta<- zibetaF(pres1)


#Hbeta is
pi%*%ziBeta


#Check that this is the same as the standard calculation of Hbeta using the vegetarian package

Hpres1<-log(d(pres1,lev="beta",wts=pj))


#change in diversity:


Hpres1-Hpast

Selection<-(pi-piPast)%*%ziBeta
Transmission<-pi%*%(ziBeta-ziBetaPast)


Selection+Transmission
Hpres1-Hpast


###############


calculateDeltaD<-function(past, pres, divOrder){
  
  pjpast<-rowSums(past)/sum(past)
  pjprest<-rowSums(pres)/sum(pres)
  
  d(pres,lev="beta",wts=pjprest,q=divOrder)-d(past,lev="beta",wts=pjpast,q=divOrder)
  print(d(pres,lev="beta",wts=pjprest,q=divOrder))
}

calculateDeltaH<-function(past,pres){
  pjpast<-rowSums(past)/sum(past)
  pjprest<-rowSums(pres)/sum(pres)
  
  #d returns jost's "true" beta diversity, loging rescales to be shannon wiener beta diveristy
  log(d(pres,lev="beta",wts=pjprest))-log(d(past,lev="beta",wts=pjpast))
}

#this is panel c in the current figure seleciton for a generalist decreases beta diversity
past<-cbind(s1=c(1,1), #abundance of species 1 in each patch
            s2=c(0,1)) #abundance of species 2 in each patch


pres<-cbind(s1=c(2,2), #abundance of species 1 in each patch
             s2=c(0,1)) #abundance of species 2 in each patch

#summary statistics needed to compute hbeta
divOrder<-1
pjpast<-rowSums(past)/sum(past)
pjprest<-rowSums(pres)/sum(pres)
log(d(pres,lev="beta",wts=pjprest,q=divOrder))-log(d(past,lev="beta",wts=pjpast,q=divOrder))


calculateDeltaD(past,pres,0)
calculateDeltaD(past,pres,1)
calculateDeltaD(past,pres,2)
calculateDeltaD(past,pres,3)

calculateDeltaH(past,pres)

#this is panel d in the current figure variable selection increases beta diversity
past<-cbind(s1=c(1,2), #abundance of species 1 in each patch
            s2=c(2,1)) #abundance of species 2 in each patch


pres<-cbind(s1=c(0,2), #abundance of species 1 in each patch
            s2=c(2,0)) #abundance of species 2 in each patch

#summary statistics needed to compute hbeta
pjpast<-rowSums(past)/sum(past)
pjprest<-rowSums(pres)/sum(pres)


calculateDeltaD(past,pres,0)
calculateDeltaD(past,pres,1)
calculateDeltaD(past,pres,2)
calculateDeltaD(past,pres,3)
calculateDeltaH(past,pres)


#This is the calculation for zibeta for palms in the present


#Verify that the multiplicative partition of shanon winer diversity works

#Here is the ratio of shannon winer beta diversity in the present to past
d(pres,lev="beta",wts=pjprest,q=divOrder)/d(past,lev="beta",wts=pjpast,q=divOrder)

#a reminder that this is equal to e raised to the power of the difference in shannon wiener entropy
exp(log(d(pres,lev="beta",wts=pjprest,q=divOrder))-log(d(past,lev="beta",wts=pjpast,q=divOrder)))

#from the main text, we know that for this example


#Re-check this
exp(Selection+Transmission)


#this is panel e in the current figure selection for a generalist increases beta diversity
past<-cbind(s1=c(1,0), #abundance of species 1 in each patch
            s2=c(0,2)) #abundance of species 2 in each patch


pres<-cbind(s1=c(2,0), #abundance of species 1 in each patch
            s2=c(0,2)) #abundance of species 2 in each patch

#summary statistics needed to compute hbeta


calculateDeltaD(past,pres,0)
calculateDeltaD(past,pres,1)
calculateDeltaD(past,pres,2)
calculateDeltaD(past,pres,3)
calculateDeltaH(past,pres)


#this is panel f in the current figure variable selection increases beta diversity
past<-cbind(s1=c(1,2), #abundance of species 1 in each patch
            s2=c(2,1)) #abundance of species 2 in each patch


pres<-cbind(s1=c(1,1), #abundance of species 1 in each patch
            s2=c(1,1)) #abundance of species 2 in each patch

#summary statistics needed to compute hbeta


calculateDeltaD(past,pres,0)
calculateDeltaD(past,pres,1)
calculateDeltaD(past,pres,2)
calculateDeltaD(past,pres,3)

calculateDeltaH(past,pres)


####################Code associated with appendix S2.3 #####################


#Figure S2.3
past<-cbind(s1=c(1,1), #abundance of species 1 in each patch
            s2=c(0,1)) #abundance of species 2 in each patch

#here is a model of just selection where species i produces more decendants than species j 
#but there is no dispersal among patches, as in Figure S1a
pres1<-past
pres1[,1]<-pres1[,1]*2

#Here dispersal has been added as in Figure S1b
pres2<-pres1
pres2[1:2,1]<-pres2[2:1,1]


#The partitions of selection and transmission are unchanged. 
divPart(past,pres1)
divPart(past,pres2)


#Figure S2
past<-cbind(s1=c(1,2), #abundance of species 1 in each patch
            s2=c(2,1)) #abundance of species 2 in each patch

#here is a model of just selection where both species fail to reproduce in the patch where they
#are rare, as in Figure S2 A.
pres1<-past
pres1[c(1,4)]<-pres1[c(1,4)]*0


#Here is a model of selection followed by dispersal as in Figure S2 B

pres2<-past

#Both species produce more descendants in the patch where they are rare
pres2[c(1,4)]<-pres2[c(1,4)]*2

#and none in the patches where they are common
pres2[2:3]<-0

#individuals disperse from one patch to the other
pres2[1:4]<-pres2[c(2:1,4:3)]


#The partitions of change in shannon wiener entropy are the same
divPart(past,pres1)
divPart(past,pres2)

#the interpretation of selection is the same in both cases,
#the frequencies of both species across the metacommunity are unchanged, i.e. selection=0.


#Transmission measures effects that are missed by the selection term, but 
#different effects can lead to the same transmission term.
