## Supplemental appendix 2 for "Disentangling niche theory and beta diversity change"

### Appendix S 2.1: beta diversity and species community divergence

For Shannon-Winer entropy, beta diversity is defined as the difference between gamma and alpha entropy:

$$(1) \qquad H_\beta = H_\gamma - H_\alpha$$

We will re-express $H_\beta$ as an average of species' contributions weighted by species' frequency:

$$(2) \qquad H_\beta = \sum_i p_{i\bullet} z_{i\beta}$$

To do this we first re-express $H_\alpha$, $H_\gamma$ in the same form, then use the difference of these two expressions to find $z_{i\beta}$.

#### Expression $H_\alpha$ as an average of species measurements

The true Shannon wiener alpha diversity is (Jost 2017; eq 11b):

$$(3) \qquad {}^1D_\alpha = exp\left[-w_1 \sum_{i=1}^{S}(p_{i1} lnp_{i1}) + -w_2 \sum_{i=1}^{S}(p_{i2} lnp_{i2}) + \cdots\right]$$

Here $p_{i1}$ is the frequency of species $i$ in community 1, $\sum_{i=1}^{S}(p_{i1} lnp_{i1})$ is the Shannon Wiener entropy of community 1. The expression in square brackets is an average of diversity among individual communities (implicitly 1…N), where each community is assigned a weight (for example $w_1$ for community 1).

For Shannon Wiener entropy, alpha diversity is found by taking the log of ${}^1D_\alpha$ (Jost 2006) giving:

$$(4) \qquad H_\alpha = -w_1 \sum_{i=1}^{S}(p_{i1} lnp_{i1}) + -w_2 \sum_{i=1}^{S}(p_{i2} lnp_{i2}) + \cdots$$

This notation can be made more compact by indexing communities j=1…N and using summation notation over communities. To maintain consistency with notation in information theory (Cover and Thomas 2012), relabel Jost's term for the probability of species *i* in a given community: $p_{i1}$ as the probability that an individual belongs to species *i* given that it occupies community *j*: $p_{i|j} = \frac{p_{ij}}{p_j}$. In this revised terminology $p_{ij}$ is the probability that an individual in the community is a member of species *i* and occupying community *j*. Substituting these definitions into equation 4 gives:

$$(5) \qquad H_\alpha = \sum_{j=1}^{N} \sum_{i=1}^{S} p_{i|j} w_j \left(-ln\left(p_{i|j}\right)\right)$$

To express equation 5 as a weighted average analogous to equation 2 substitute in $\frac{p_i}{p_i} = 1$ giving:

$$(6) \qquad H_\alpha = \sum_{i=1}^{S} \frac{p_{i\bullet}}{p_{i\bullet}} \sum_{j=1}^{N} p_{i|j} w_j \left(-ln\left(p_{i|j}\right)\right)$$

$$(7) \qquad H_\alpha = \sum_{i=1}^{S} p_{i\bullet} \sum_{j=1}^{N} \frac{p_{i|j}}{p_{i\bullet}} w_j \left(-ln\left(p_{i|j}\right)\right)$$

$$(8) \qquad H_\alpha = \sum_{i=1}^{S} p_{i\bullet} \sum_{j=1}^{N} \frac{p_{ij} w_j}{p_{\bullet j} p_{i\bullet}} \left(-ln\left(p_{i|j}\right)\right)$$

This equation expresses alpha entropy as an average of each species' measurement: $z_{i\alpha,w} = \sum_{j=1}^{N} \frac{p_{ij} w_j}{p_{\bullet j} p_{i\bullet}} \left(-ln\left(p_{i|j}\right)\right)$, weighted by species' frequency. Note that $z_{i\alpha,w}$ includes a weighting term $w_j$. In the main text we simplify this further by weighting all communities by the number of individual they contain, $w_j = p_{\bullet j}$. When this is done, equation 8 reduces to:

$$(9) \; H_\alpha = \sum_{i=1}^{S} p_{i\bullet} \sum_{j=1}^{N} p_{j|i} \left(-ln\left(p_{i|j}\right)\right)$$

This can be re-expressed in the same form as equation 2 by defining a new variable: $z_{i\alpha} = \sum_{j=1}^{N} p_{j|i} \left(-ln\left(p_{i|j}\right)\right)$ the average rarity of species i across all communities, weighted by the frequency of each community.

Gamma entropy can be expressed as:

(10) $H_\gamma = \sum_i p_{i\bullet}(-log(p_{i\bullet}))$

This means that the measurement of interest for gamma entropy is: $z_{i\gamma} = -log(p_{i\bullet})$.

### Expression for $H_\beta$ as an average of species measurements

Since:

(11) $H_\beta = H_\gamma - H_\alpha$

It follows that

(12) $H_\beta = \sum_i p_{i\bullet} z_{i\gamma} - \sum_i p_i z_{i\alpha}$

(13) $H_\beta = \sum_i p_{i\bullet}(z_{i\gamma} - z_{i\alpha})$

So the measurement of interest for $H_\beta$ is:

(14) $z_{i\beta} = z_{i\gamma} - z_{i\alpha}$

By combining equation 14 with the definitions of $z_{i\gamma}$ and $z_{i\alpha}$ we obtain:

(15) $z_{i\beta} = -log(p_{i\bullet}) - \sum_{j=1}^{N} p_{j|i}\left(-ln(p_{i|j})\right)$

$$= \sum_{j=1}^{N} p_{j|i}(-log(p_{i\bullet})) - \sum_{j=1}^{N} p_{j|i}\left(-ln(p_{i|j})\right)$$

$$= \sum_j p_{j|i}\left((-log(p_{i\bullet})) - \left(-ln(p_{i|j})\right)\right)$$

$$= \sum_j p_{j|i}\left(-log\left(\frac{p_{i\bullet}}{p_{i|j}}\right)\right)$$

Alternatively, we can obtain an expression for species' contributions to beta entropy that has another weighting by substituting $z_{i\alpha} = z_{i\alpha,w}$ into equation 14 giving:

$$(16) \qquad z_{i\beta,w} = -log(p_{i\bullet}) - \sum_{j=1}^{N} \frac{p_{ij}w_j}{p_{\bullet j}p_{i\bullet}}\left(-ln\left(p_{i|j}\right)\right).$$

Ecologists commonly weight all communities equally which can be done by setting $w_j = \frac{1}{N}$.

### Appendix S 2.2: Partitioning change in other beta diversity metrics

In this section we outline the steps needed to partition selection and transmission for measures of beta diversity other than Shannon Wiener entropy. A major upshot of this proof is that a general expression for beta diversity change will be possible but difficult to intuit. Jost 2007's suggested generalization of beta diversity is the "true" gamma diversity divided by "true" alpha diversity:

$$(17) \qquad {}^{q}D_{\beta} = \frac{{}^{q}D_{\gamma}}{{}^{q}D_{\alpha}}$$

Where (using our notation) the numbers equivalent of gamma diversity is:

$$(18) \qquad {}^{q}D_{\gamma} = \left[\sum_{i=1}^{S} p_{i\bullet}{}^{q}\right]^{\frac{1}{1-q}}$$

And the numbers equivalent of alpha diversity is:

$$(19) \qquad {}^{q}D_{\alpha} = \left[\sum_{i=1}^{S} \sum_{j=1} \frac{w_j^q p_{i|j}^q}{\sum_j w_j^q}\right]^{\frac{1}{1-q}}$$

Equation 19 can be analysed using an extension of a Price equation by defining two measurements, one to describe a species' contribution to gamma diversity ($z_{i\gamma q} = p_{i\bullet}{}^{q-1}$) and one to describe their contribution to alpha diversity and a measurement associated with alpha diversity ($z_{i,\alpha,q}$). This is found by multiplying Equation 20 by $\frac{p_{i\bullet}}{p_{i\bullet}} = 1$ giving:

$$(20) \qquad =\left[\sum_{i=1}^{S} \frac{p_{i\bullet}}{p_{i\bullet}} \sum_{j=1} \frac{w_j^q p_{i|j}^q}{\sum_j w_j^q}\right]^{\frac{1}{1-q}}$$

$$=\left[\sum_{i=1}^{S} p_{i\bullet} \sum_{j=1} \frac{p_{i|j}^q}{p_{i\bullet}} \frac{w_j^q}{\sum_j w_j^q}\right]^{\frac{1}{1-q}}$$

$$=\left[\sum_{i=1}^{S} p_{i\bullet} z_{i,\alpha,q}\right]^{\frac{1}{1-q}}$$

Where $z_{i,\alpha,q} = \sum_{j=1} \frac{p_{i|j}^q}{p_{i\bullet}} \frac{w_j^q}{\sum_j w_j^q}$

Beta diversity is, ${}^qD_\beta$ is:

$$(21) \qquad {}^qD_\beta = \left(\frac{\sum_{i=1}^{S} p_i z_{i,\gamma,q}}{\sum_{i=1}^{S} p_i z_{i,\alpha,q}}\right)^{\frac{1}{1-q}}$$

To partition change in ${}^qD_\beta$, it will help to simplify notation by defining:

$$(22) \qquad {}^qD_\beta = f(p_i, z_{i,\gamma,q}, z_{i,\alpha,q})$$

This function can be partitioned using the method presented in (Frank and Godsoe 2020) to produce selection and transmission terms:

$$(23) \qquad \Delta f(p_i, z_{i,\gamma,q}, z_{i,\alpha,q}) = \underbrace{f(p_i', z_{i,\gamma,q}, z_{i,\alpha,q}) - f(p_i, z_{i,\gamma,q}, z_{i,\alpha,q})}_{selection/drift} +$$

$$\underbrace{f(p_i', z_{i,\gamma,q}', z_{i,\alpha,q}') - f(p_i', z_{i,\gamma,q}, z_{i,\alpha,q})}_{Transmission}$$

This expression shows that partitioning of beta diversity into selection and transmission is possible, but it is a non-linear function of multiple traits. We leave it to the reader to see if the insights from this expression justify its additional complexity over the expressions derived in the main text.

This partitioning can be illustrated with special cases, one that is of particular interest is species richness. We present an analysis of there but will echo a point we made in (Godsoe et al. In press), selection is an imperfect tool to study species richness because its effects are often exactly opposed by transmission. To derive an expression for beta diversity in this case set q=0 producing:

$$z_{i,\gamma,0} = p_{i\bullet}^{0-1} \tag{24}$$

$$= \frac{1}{p_i}.$$

And:

$$z_{i,\alpha,0} = \sum_{j=1} \frac{p_{i|j}^0}{p_{i\bullet}} \frac{w_j^0}{\sum_j w_j^0}$$

$$z_{i,\alpha,0} = \frac{1}{Jp_{i\bullet}} \sum_{j=1} p_{i|j}^0$$

Under the convention that $0^0 = 0$, $\sum_{j=1} p_{i|j}^0 / N$ is the proportion of communities occupied by species *i*, hereafter $p_{i\bullet,occ}$. So

$$z_{i,\alpha,0} = \frac{p_{i\bullet,occ}}{p_{i\bullet}}$$

This is the ratio of the frequency of communities occupied by a species to the frequency of individuals belonging to that species. It exceeds 1 when the species occupies many communities, relative to its frequency in the metacommunity. It is less than 1 when the species occupies few communities relative to its frequency. An artificial example can help to illustrate this trait. Consider a metacommunity consisting of 9 pigeons in community 1 and 1 wren in community 2. Each species occupies ½ of the available communities but they have different scores for $z_{i,\alpha,0}$ because the vast majority of individuals are pigeons and a small minority are wrens. Pigeons have a small score because they occupy relatively few communities compared with their frequency $z_{i,\alpha,0} = \frac{1/2}{9/10} = \frac{5}{9}$. In contrast wrens have a high score because relative the number of communities they occupy their frequency is quite low when $z_{i,\alpha,0} = \frac{1/2}{1/10} = 5$.

In a metacommunity where species frequencies change but no species occupy new communities or become extirpated selection and transmission balance out completely. In other words total change is:

(25) $\Delta\ {}^1D_\beta =$

$$\underbrace{\frac{\sum_{i=1}^{S} \frac{p_{i\bullet}'}{p_{i\bullet}}}{\sum_{i=1}^{S} \frac{p_{i\bullet}' p_{i\bullet,occ}}{p_{i\bullet}}} - \frac{\sum_{i=1}^{S} \frac{p_{i\bullet} 1}{p_{i\bullet}}}{\sum_{i=1}^{S} \frac{p_{i\bullet} p_{i\bullet,occ}}{p_{i\bullet}}}}_{Selection+drift} + \underbrace{\frac{\sum_{i=1}^{S} \frac{p_{i\bullet}'}{p_{i\bullet}'}}{\sum_{i=1}^{S} \frac{p_i' p_{i\bullet,occ}'}{p_{i\bullet}'}} - \frac{\sum_{i=1}^{S} \frac{p_{i\bullet}'}{p_{i\bullet}}}{\sum_{i=1}^{S} \frac{p_{i\bullet}' p_{i\bullet,occ}}{p_{i\bullet}}}}_{Transmission}$$

This can be simplified to:

(26) $$\Delta\ {}^1D_\beta = \frac{\sum_{i=1}^{S} \frac{p_{i\bullet}'}{p_{i\bullet}}}{\sum_{i=1}^{S} \frac{p_{i\bullet}'}{p_{i\bullet}} p_{i,occu}} - \frac{S}{\sum_{i=1}^{S} p_{i\bullet,occ}} + \frac{S}{\sum_{i=1}^{S} p_{i\bullet,occ}'} - \frac{\sum_{i=1}^{S} \frac{p_{i\bullet}'}{p_{i\bullet}}}{\sum_{i=1}^{S} \frac{p_{i\bullet}'}{p_{i\bullet}} p_{i\bullet,occ}}$$

Since the number of communities occupied remains constant: $p_{i\bullet,occupied}' = p_{i\bullet,occupied}$. When this substitution is made in to Equation 26 $\Delta\ {}^1D_\beta = 0$ .

To conclude, in the case of beta diveristy based on species richness, the effect of selection is frequently obscured by transmission. This suggests that an analysis of selection for this particular diveristy metric is of limited utility.

### Appendix S 2.3: New arrivals to the metacommunity

To allow immigrants from outside of the metacommunity we can use an extension of the Price equation (Kerr and Godfrey-Smith 2009, Frank 2012). This approach distinguishes two categories of individuals: 1) individuals who descended from members of the past community (the frequency of such individuals is $\omega$), 2) individuals that are not descended from past members of the community, such as immigrants (the frequency of such individuals is $\mu$). Individuals in the present community are either descendants or immigrants such that $\omega + \mu = 1$. This leads to the following formula for $H_\beta$ in the present community:

(27)
$$D' = \omega \sum_i p_{i\bullet|D} z_{i\beta}{}' + \mu \sum_k p_{k\bullet|I} z^*_{k\beta}$$

$p_{i\bullet|D}$ represents the frequency of species *i* among individuals in the present community descended from individuals that were present in the past observation period, $p_{k\bullet|I}$ is the frequency of immigrants belonging to species *k* among all immigrants. Thus $z_{i\beta}{}'$ is the divergence of the $i^{\text{th}}$ descendant species in the present community. $z^*_{k\beta}$ is the divergence of the $k^{\text{th}}$ immigrant species in the present community. Individuals that are descendants of the past community are indexed separately from immigrants, even when both belong to the same species.

Change in beta diversity can now be partitioned using the "Extended Price equation" (Kerr and Godfrey-Smith 2009, Frank 2012):

(28)
$$\Delta H_\beta = \underbrace{\omega \sum_i (p_{i\bullet|D} - p_{i\bullet}) z_{i\beta}}_{\substack{Selection\ and \\ drift}} + \underbrace{\omega \sum_i p_{i\bullet|D}{}' \Delta z_{i\beta}}_{\substack{change \\ in\ z_{i\beta}}} + \underbrace{\mu \sum_k p_{k\bullet|I} z^*_{k\beta} - \mu \sum_i p_{i\bullet} z_{i\beta}}_{immigration}$$

Much like in Equation 4 in the main text, descendants' contributions to $H_\beta$ change are partitioned into selection and change in divergence. The effect of these mechanisms is weighted by the frequency of descendants in the present community $\omega$. The effects of immigration depend on the proportion of individuals in the present community that are immigrants ($\mu$), and the difference between the average divergence of immigrants in the present, and the average divergence of the past community. The immigration term is small when immigrants are rare (i.e. when $\mu$ is close to 0) and the immigration term is small when the diversity of immigrants is similar to the diversity of ancestors (i.e. when $\sum_k p_{k\bullet|I} z^*_{k\beta} - \sum_i p_{i\bullet} z_{i\beta}$ is small. To find equivalent expressions for alpha or gamma diversity change, replace the $\beta$ subscript with the subscript of the desired diversity component.

#### Appendix S 2.4: Immigration within the metacommunity

In the main text we state that dispersal within the metacommunity does not change the partitioning of selection and changes in $z_{i\beta}$. This is illustrated in figure S1. Figure S2 illustrates how dispersal can change the interpretation of changes in $z_{i\beta}$.

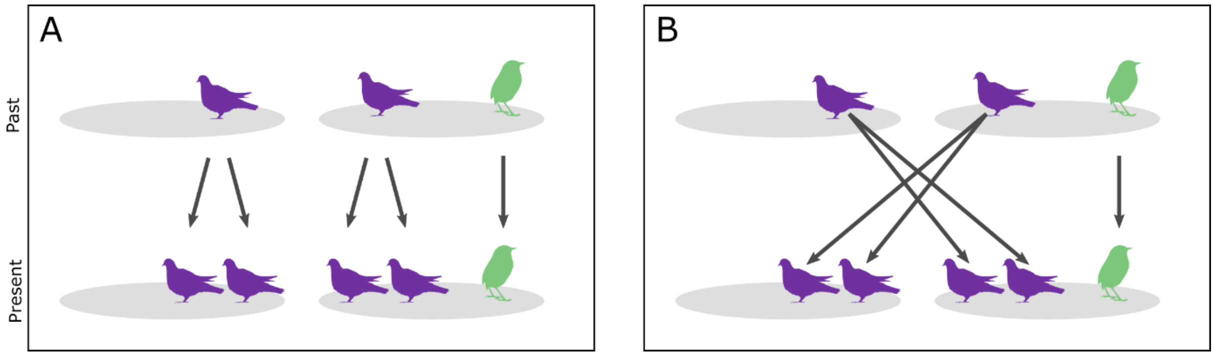


Figure S1 Illustrates why the partitioning of selection and divergence change is insensitive to immigration within the metacommunity. A) shows a scenario with selection across the metacommunity but no immigration (i.e. all descendants reside in the same community as their ancestors). B) shows a scenario with comparable selection across the metacommunity and immigration (i.e. the descendant pigeons live in a different community from their ancestors). In both A and B the initial conditions are the same and the total number of descendants of each species in each community are the same. This is enough to guarantee that the total diversity change and the partitioning between selection and change in divergence is identical. See R script in Appendix S1 for example calculations.

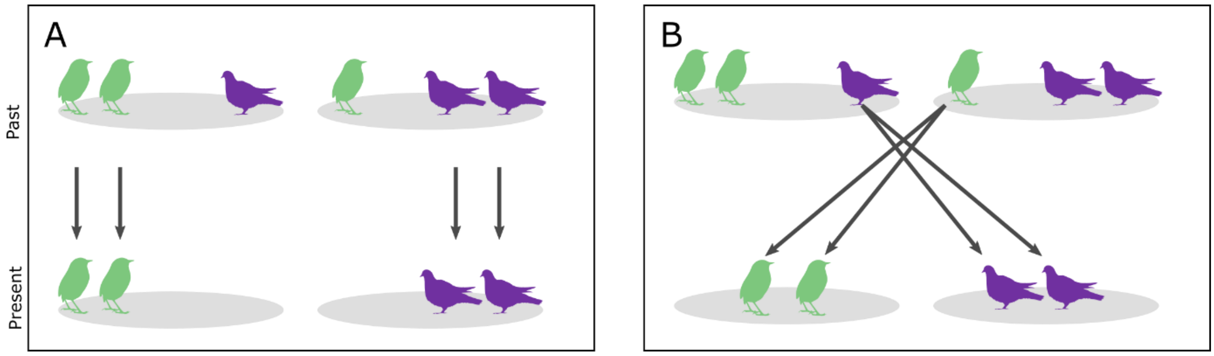


Figure S2 Illustrates how information on immigration can help to clarify the interpretation of the divergence change term. A) Shows a scenario with spatially variable selection across the metacommunity but no immigration (i.e. all descendants reside in the same community as their ancestors). B) Shows a scenario with spatially variable selection and immigration (i.e. some descendant birds moved to another community). In both scenarios the initial conditions are the same and the total number of descendants of each species in each community are the same. This is enough to guarantee that the beta diversity change and its partitions remain the same. But the average change in $z_{i\beta}$ has a different interpretation in each panel. In A the average change in $z_{i\beta}$ represents spatially variable selection that favours each species when it is common. B, the average change in $z_{i\beta}$ is a consequence of two different forces, spatially variable selection which favours each species when it is rare and immigration.
