## Supplementary material for "Disentangling niche theory and beta diversity change": example code folder: bootstrap2 may 2019.R

```
#Code for Godsoe et al 2021 Disentangling niche theory and beta diversity change

#Boostrapping code was developed in R v 3.6.1
#Code was last run in 4.0.2


#21 05 2019 this code is run by analysis for draft 1.rmd

#22/03/2019
#Will tried to adopt the code for his examples


# 21/03/2019
# by Elena Moltchanova
# making sense of Will's diversity functions :)
#rm(list=ls())
#setwd("H\\collabs\\Will\\Diversity\\")
#source("H:\\collabs\\Will\\Diversity\\will_function.txt")

#source("prelim analysis.R")


#d84 <- read.csv("data1984.csv")
#d94 <- read.csv("data1994.csv")

past <- data.matrix(d84)
present <- data.matrix(d94)

divSummaries<-data.frame(selection=NA,immigration=NA,transmission=NA)
ppSummary<-propPieceWImmigrants(past=past,pres=present)

################partition  diversity############ 

#gamma diversity
divSummaries[1,1:3]<-classicPriceExtended1(ppSummary$past$pi,       #frequency of ancestors
                                           ppSummary$pres$pidecendants,      #frequency individuals decendend from ancestral community
                                           ppSummary$pres$freqofImmigrantpool,      #frequency of each unconnected type
                                           ppSummary$past$Zgamma,         #traits of ancestors
                                           ppSummary$pres$Zgamma,         #traits of decendants community
                                           ppSummary$pres$ZgammaImm,       #trait value of unconnected populations
                                           ppSummary$pres$pInImmigrantpool)


#alpha diversity
divSummaries[2,1:3]<-classicPriceExtended1(ppSummary$past$pi,       #frequency of ancestors
                                           ppSummary$pres$pidecendants,      #frequency individuals decendend from ancestral community
                                           ppSummary$pres$freqofImmigrantpool,      #frequency of each unconnected type
                                           ppSummary$past$Zalpha,         #traits of ancestors
                                           ppSummary$pres$Zalpha,         #traits of decendants community
                                           ppSummary$pres$ZalphaImm,       #trait value of unconnected populations
                                           ppSummary$pres$pInImmigrantpool) 

#beta diversity
divSummaries[3,1:3]<-classicPriceExtended1(ppSummary$past$pi,       #frequency of ancestors
                                           ppSummary$pres$pidecendants,      #frequency individuals decendend from ancestral community
                                           ppSummary$pres$freqofImmigrantpool,      #frequency of each unconnected type
                                           ppSummary$past$Zbeta,         #traits of ancestors
                                           ppSummary$pres$Zbeta,         #traits of decendants community
                                           ppSummary$pres$ZbetaImm,       #trait value of unconnected populations
                                           ppSummary$pres$pInImmigrantpool) 

divSummaries
rowSums(divSummaries)


### let's try bootstrap
#this construts a 3d array of dimensions 3 X 4 X nboot. 
Nboot <- 10^4
div.boot <- array(dim=c(3,4,Nboot))


for(b in 1:Nboot){print(b)
    #Create a new array of abunances. 
    #To get this, Use the poisson distribution to simulate counts for each element of the present (i.e. 4x71) 
    #Observed counts for the present are used. 


  past.sim <- past#array(rpois(prod(dim(past)),past),dim=dim(past))
  present.sim <- array(rpois(prod(dim(past)),present),dim=dim(present))


  ppSummary<-propPieceWImmigrants(past=past.sim,pres=present.sim)
  
  #gamma diversity
  div.boot[1,1:3,b]<-classicPriceExtended1(ppSummary$past$pi,       #frequency of ancestors
                                             ppSummary$pres$pidecendants,      #frequency individuals decendend from ancestral community
                                             ppSummary$pres$freqofImmigrantpool,      #frequency of each unconnected type
                                             ppSummary$past$Zgamma,         #traits of ancestors
                                             ppSummary$pres$Zgamma,         #traits of decendants community
                                             ppSummary$pres$ZgammaImm,       #trait value of unconnected populations
                                             ppSummary$pres$pInImmigrantpool)
  
  #alpha diversity
  div.boot[2,1:3,b]<-classicPriceExtended1(ppSummary$past$pi,       #frequency of ancestors
                                             ppSummary$pres$pidecendants,      #frequency individuals decendend from ancestral community
                                             ppSummary$pres$freqofImmigrantpool,      #frequency of each unconnected type
                                             ppSummary$past$Zalpha,         #traits of ancestors
                                             ppSummary$pres$Zalpha,         #traits of decendants community
                                             ppSummary$pres$ZalphaImm,       #trait value of unconnected populations
                                             ppSummary$pres$pInImmigrantpool) 
  
  #beta diversity
  div.boot[3,1:3,b]<-classicPriceExtended1(ppSummary$past$pi,       #frequency of ancestors
                                             ppSummary$pres$pidecendants,      #frequency individuals decendend from ancestral community
                                             ppSummary$pres$freqofImmigrantpool,      #frequency of each unconnected type
                                             ppSummary$past$Zbeta,         #traits of ancestors
                                             ppSummary$pres$Zbeta,         #traits of decendants community
                                             ppSummary$pres$ZbetaImm,       #trait value of unconnected populations
                                             ppSummary$pres$pInImmigrantpool) 
  
  #this computes the total change in diversity
  div.boot[,4,b] <- rowSums(div.boot[,1:3,b])
} # end of b

### SUMMARIES
div.boot.lo <- div.boot.hi <- div.boot.se <- div.boot.mn <- div.boot.p <- array(dim=c(3,4))
for(i in 1:3){
for(j in 1:4){
div.boot.lo[i,j] <- quantile(div.boot[i,j,],.025)
div.boot.hi[i,j] <- quantile(div.boot[i,j,],.975)
div.boot.se[i,j] <- sd(div.boot[i,j,])
div.boot.mn[i,j] <- mean(div.boot[i,j,])
div.boot.p[i,j] <- mean(div.boot[i,j,]>0)
}}

div.boot.lo
div.boot.hi
div.boot.se
div.boot.mn
div.boot.p


# Was gamma diversity most influenced by immigration, selection or transmission?
table(apply(div.boot[1,1:3,],2,which.max))/Nboot

#i think i should read this as out of all simulations 93.85% had selection 
#stronger than immigration, 6.15 % had immigration stronger than selection
#in none of the simulations was transmission the strongst mechanism. 

# Did selection most increase alpha diversity or beta diversity
#this hypothesis test is not ideal because change in beta diversity is negative,
mean(div.boot[1,2,]>div.boot[1,3,])

#div.boot.lo<-round(div.boot.lo,3)
#div.boot.hi<-round(div.boot.hi,3)
matrix(paste(div.boot.lo,div.boot.hi,sep=""),3,4)


#Ok, let's try to convey the results as a figure. 
```
