## Supplementary material for "Disentangling niche theory and beta diversity change": example code folder: bootstrap drift test oct 19.R

```
#Code for Godsoe et al 2021 Disentangling niche theory and beta diversity change

#Boostrapping code was developed in R v 3.6.1
#Code was last run in 4.0.2


#########################code duplicated from bootstrap2 may 2019 to get observed values########################


divSummaries<-data.frame(selection=NA,immigration=NA,transmission=NA)
ppSummary<-propPieceWImmigrants(past=past,pres=present)

################partition  diversity############ 

#gamma diversity
divSummaries[1,1:3]<-classicPriceExtended1(ppSummary$past$pi,       #frequency of ancestors
                                           ppSummary$pres$pidecendants,      #frequency individuals decendend from ancestral community
                                           ppSummary$pres$freqofImmigrantpool,      #frequency of each unconnected type
                                           ppSummary$past$Zgamma,         #traits of ancestors
                                           ppSummary$pres$Zgamma,         #traits of decendants community
                                           ppSummary$pres$ZgammaImm,       #trait value of unconnected populations
                                           ppSummary$pres$pInImmigrantpool)


#alpha diversity
divSummaries[2,1:3]<-classicPriceExtended1(ppSummary$past$pi,       #frequency of ancestors
                                           ppSummary$pres$pidecendants,      #frequency individuals decendend from ancestral community
                                           ppSummary$pres$freqofImmigrantpool,      #frequency of each unconnected type
                                           ppSummary$past$Zalpha,         #traits of ancestors
                                           ppSummary$pres$Zalpha,         #traits of decendants community
                                           ppSummary$pres$ZalphaImm,       #trait value of unconnected populations
                                           ppSummary$pres$pInImmigrantpool) 

#beta diversity
divSummaries[3,1:3]<-classicPriceExtended1(ppSummary$past$pi,       #frequency of ancestors
                                           ppSummary$pres$pidecendants,      #frequency individuals decendend from ancestral community
                                           ppSummary$pres$freqofImmigrantpool,      #frequency of each unconnected type
                                           ppSummary$past$Zbeta,         #traits of ancestors
                                           ppSummary$pres$Zbeta,         #traits of decendants community
                                           ppSummary$pres$ZbetaImm,       #trait value of unconnected populations
                                           ppSummary$pres$pInImmigrantpool) 


##########################this is the test for whether lineage sorting can be explained by drift#####################


Nboot <- 10^4
div.boot2 <- array(dim=c(3,4,Nboot))

for(b in 1:Nboot){#print(b)
  
  # ASSUMPTIONS
  # 1. past is fixed
  # 2. present counts are proportional to past counts (meaning no immigration)
  # 3. expected present community total is the observed present (descendants) community total
  
  # checking which species were present ANYWHERE in the past
  is.descendant <- (apply(past,2,sum)>0)
  
  # total present number of individuals belonging to descendant species in each location
  present.descendants <- c(present%*%is.descendant)
  
  # total past number of individuals belonging to descendant species in each location 
  past.descendants <- c(past%*%is.descendant)
  
  # past if fixed
  past.sim <- past
  
  # present  #create simulations of present counts where the abundance of each species 
  present.sim <-  array(rpois(prod(dim(present)), #the array will include species*patch entries
                              past*present.descendants/past.descendants), #the lambda parameter is equal to the abundance in the past times the change in community size.
                        dim=dim(present))
  
  
  ppSummary<-propPieceWImmigrants(past=past,pres=present.sim)
  
  #gamma diversity
  div.boot2[1,1:3,b]<-classicPriceExtended1(ppSummary$past$pi,       #frequency of ancestors
                                            ppSummary$pres$pidecendants,      #frequency individuals decendend from ancestral community
                                            ppSummary$pres$freqofImmigrantpool,      #frequency of each unconnected type
                                            ppSummary$past$Zgamma,         #traits of ancestors
                                            ppSummary$pres$Zgamma,         #traits of decendants community
                                            ppSummary$pres$ZgammaImm,       #trait value of unconnected populations
                                            ppSummary$pres$pInImmigrantpool)
  
  #alpha diversity
  div.boot2[2,1:3,b]<-classicPriceExtended1(ppSummary$past$pi,       #frequency of ancestors
                                            ppSummary$pres$pidecendants,      #frequency individuals decendend from ancestral community
                                            ppSummary$pres$freqofImmigrantpool,      #frequency of each unconnected type
                                            ppSummary$past$Zalpha,         #traits of ancestors
                                            ppSummary$pres$Zalpha,         #traits of decendants community
                                            ppSummary$pres$ZalphaImm,       #trait value of unconnected populations
                                            ppSummary$pres$pInImmigrantpool) 
  
  #beta diversity
  div.boot2[3,1:3,b]<-classicPriceExtended1(ppSummary$past$pi,       #frequency of ancestors
                                            ppSummary$pres$pidecendants,      #frequency individuals decendend from ancestral community
                                            ppSummary$pres$freqofImmigrantpool,      #frequency of each unconnected type
                                            ppSummary$past$Zbeta,         #traits of ancestors
                                            ppSummary$pres$Zbeta,         #traits of decendants community
                                            ppSummary$pres$ZbetaImm,       #trait value of unconnected populations
                                            ppSummary$pres$pInImmigrantpool) 
  
  div.boot2[,4,b] <- rowSums(div.boot2[,1:3,b])
} # end of b

divobs <- cbind(divSummaries, rowSums(divSummaries)) # this should be the observed selection term

div.boot.lo2 <- div.boot.hi2 <- div.boot.mn2 <- div.boot.p2 <- array(dim=c(3,4))

for(i in 1:3){                                          #for three diversity components
  for(j in 1:4){                                        #compute for four components (selection, transmission, immigration, total)
    div.boot.lo2[i,j] <- quantile(div.boot2[i,j,],.025)
    div.boot.hi2[i,j] <- quantile(div.boot2[i,j,],.975)
    div.boot.mn2[i,j] <- mean(div.boot2[i,j,])
    
    div.boot.p2[i,j] <- mean(abs(div.boot2[i,j,])>divobs[i,j])
  }}

div.boot.lo2   #these are the low values from bootstrapping
div.boot.hi2   #these are the high values from bootstrapping
div.boot.mn2   #these are the mean values from bootrapping
divobs

div.boot.p2   #in how many simulations is the absolute value of the bootstrap lineage sorting greater than that expected by drift


#draw a histogram of all values for alpha diversity


#correction by WG so the price equation "selection term" is the strength of selection (sum deltapi *zi) times omega (thefrequency of decendants)
#this part of code converts this back into the "pure" strength of selection i.e. 


omega<-sum(present.descendants)/sum(present)
ObsLSCorrected<-divobs$selection/omega

pvalues<-vector(mode="numeric",3)
for(i in 1:3){
  pvalues[i]<-mean(ObsLSCorrected[i]<div.boot2[i,1,])
  
}

#I don't think the paper needs a histogram illustration of the test, but i've cooked one up 
#for my own sake. 
#hist(div.boot2[1,1,])
#points(SDpiZi[1],0,pch=16, col="red", cex=3)
#abline(v=div.boot.hi2[1,1], col="blue")
#abline(v=div.boot.lo2[1,1], col="blue")


driftTest<-data.frame(lowerCi=div.boot.lo2[,1], upperCI=div.boot.hi2[,1], ObsLSCorrected=ObsLSCorrected,pvalue=pvalues)

#row.names(driftTest)<-c("Gamma", "Alpha", "Beta")
```
