## Supplementary material for "Disentangling niche theory and beta diversity change": example code folder: reproducible analyses.R

```
#Code for Godsoe et al 2021 Disentangling niche theory and beta diversity change

#Boostrapping code was developed in R v 3.6.1
#Code was last run in 4.0.2


#replicate analyses for paper

#the analyses presented in the draft manuscript as of august 2020
#are based on the rmd file "analyses for draft7" however
#these use R scripts based on a version of hte dataset including data that is unpublished and collected by 
#edmund tanner, since this is not data associated with the paper we are presenting a revised version
# here. In principle this script gives the same answers (we have done several unit checks), the figures look similar 
#but lack some of the formatting commands in Rmarkdown.


require(RColorBrewer)
require(ggplot2)


#Contact william godsoe for queries
#to run this code you will need two RData files TannerTable84.RData, and TannerTable94.RData
#These contain tables of counts for each species at each of four sites
#in additoin you will require two additional scripts that are used to run simulation analysis
#"bootstrap2 may 2019.R" and "bootstrap drift test oct 19.R"

rm(list=ls())


logMod<-function(X){
  Y<-log(X)
  Y[X==0]<-0
  Y[is.infinite(Y)]<-0
  return(Y)
}

#this calculates the measurement associated with species relevant for alpha diversity
ZCalc<-function(PJGI,ZI){
  colSums(PJGI*(-logMod(ZI)),na.rm=T)
  
}


probPieces<-function(counts){
  #proportion of metacommunity in species i in patch j
  pij<-counts/sum(counts)
  
  #proportion of individuals in patch j
  pj<-rowSums(pij)
  
  #proportion of individuals in species i
  pi<-colSums(pij)
  
  #for some calculations it is easier to have a site*species matrix of pi rather
  #than a vaector
  piAll<-matrix(pi, byrow=T,nrow=dim(pij)[1],ncol=dim(pij)[2])
  
  #proportion of individuals in species i given that we are in patch j
  piGivenj<-pij/pj
  
  pjGiveni<-t(t(pij)/pi)
  
  
  Zalpha<-ZCalc(pjGiveni,piGivenj)
  Zbeta <-ZCalc(pjGiveni,piAll/piGivenj)
  Zgamma<-ZCalc(pjGiveni,piAll)
  
  return(list(pij=pij,
              pi=pi,
              pj=pj, 
              piAll=piAll,
              piGivenj=piGivenj,
              pjGiveni=pjGiveni,
              Zalpha=Zalpha,
              Zbeta=Zbeta,
              Zgamma=Zgamma))
  
}


propPieceWImmigrants<-function(past,pres){
  
  #this function accepts tables of species' counts from the past and present
  #then computes summary statistics needed to use the extended price equation with immigration
  occuredInPast<-colSums(past)>0
  immigrantSpecies<- !occuredInPast & colSums(pres)>0
  
  
  decendantsInd<-sum(pres[,occuredInPast])
  immigrantsInd<-sum(pres[,immigrantSpecies])
  totalInd<-sum(pres)
  
  
  
  #compute summaries only for species present in both datasets
  psPast<-probPieces(past[,occuredInPast])
  psPres<-probPieces(pres[,occuredInPast])
  
  psPast1<-probPieces(past)
  psPres1<-probPieces(pres)
  
  freqofImmigrantpool<-psPres1$pi[immigrantSpecies]/sum(psPres1$pi[immigrantSpecies])
  
  #this is a crude way to normalize the scores so that decendant scores sum up to 1,
  #really it would be safer to create separate vectors for decendants and immigrants, can try that later.
  psPres1$pidecendants<-psPres1$pi/sum(psPres1$pi[occuredInPast])
  psPres1$pidecendants[immigrantSpecies]<-0
  
  psPres1$freqofImmigrantpool<-psPres1$pi[immigrantSpecies]/sum(psPres1$pi[immigrantSpecies])
  psPres1$pInImmigrantpool<-immigrantsInd/totalInd
  
  psPres1$ZalphaImm<-psPres1$Zalpha[immigrantSpecies]
  psPres1$ZbetaImm<- psPres1$Zbeta[immigrantSpecies]
  psPres1$ZgammaImm<-psPres1$Zgamma[immigrantSpecies]
  
  list(past=psPast1,pres=psPres1)
}


bootStrapFun<-function(d84,d94){
  #this function calls elena's bootstrapping program then computes estimates for the importance of 
  #each component of diversity change and the importance of each mechanism, with confidence intervals
  #it requires d84 (past abundances) and d94 present abundances
  
  source("bootstrap2 may 2019.R")
  
  
  #########################this version seems to work!#################################
  
  
  
  condition=factor(c("gamma","gamma","gamma","gamma","alpha","alpha","alpha","alpha","beta","beta","beta","beta"), levels=c("gamma","alpha","beta"))
  mechanism=factor(rep(c("selection", "immigration","transmission", "total") , 3 ), levels=c("selection" , "immigration" , "transmission", "total"))
  value=as.vector(t(div.boot.mn))
  ci.lo=as.vector(t(div.boot.lo))
  ci.hi=as.vector(t(div.boot.hi))
  data=data.frame(condition,mechanism,value,ci.lo,ci.hi)
  
  
  #svg("total plot.svg")
  # Faceting
  
  
  return(data)
  
}

classicPriceExtended1 <- function(qa,qd,alpha1,za,zd,zStar,P) {
  
  #frequency in ancestral pop
  #frequency of decendants of ancestral pop
  #frequency of immigrants in species j
  #z trait value ancestors
  #trait value of decendants
  #trait value of immigrants
  #proportion of immigrants
  
  #first reformat to calculate trait change with "old species" by weeding out species only found among
  #decendants
  qaC<-qa[qa>0]
  qdC<-qd[qa>0]
  zaC<-za[qa>0]
  
  #first calculate terms involving non-immigrants
  wi=qdC/qaC                                  
  deltaZ<-zd-za
  
  
  #ignore "new species"
  deltaZ<-deltaZ[qa>0]
  
  
  term1<-sum(qaC*(wi-1)*zaC)*(1-P)
  term3<-sum(qaC*wi*(deltaZ))*(1-P)
  
  if(anyNA(alpha1)==FALSE){
    #since qd=qa(wi/wbar)
    #and wbar should be 1 in a constant population
    zBar<-sum(qa*za) #average trait value in ancestral population
    ZBarSt<-sum(alpha1*zStar)
    term2<-P*(ZBarSt-zBar)
  }
  else{
    term2<-0
  }
  
  pTerms<-c(term1,term2,term3)
  
  return(pTerms)
}


classicPriceBySpecies<-function(qa,qd,alpha1,za,zd,zStar,P) {
  
  #frequency in ancestral pop
  #frequency of decendants of ancestral pop
  #frequency of immigrants in species j
  #z trait value ancestors
  #trait value of decendants
  #trait value of immigrants
  #proportion of immigrants
  
  #first reformat to calculate trait change with "old species" by weeding out species only found among
  #decendants
  qaC<-qa[qa>0]
  qdC<-qd[qa>0]
  zaC<-za[qa>0]
  
  #first calculate terms involving non-immigrants
  wi=qdC/qaC                                  
  deltaZ<-zd-za
  
  
  #ignore "new species"
  deltaZ<-deltaZ[qa>0]
  
  
  term1<-(qaC*(wi-1)*zaC)*(1-P)
  term3<-(qaC*wi*(deltaZ))*(1-P)
  
  if(anyNA(alpha1)==FALSE){
    #since qd=qa(wi/wbar)
    #and wbar should be 1 in a constant population
    zBar<-sum(qa*za) #average trait value in ancestral population
    term2<-P*(alpha1*zStar-zBar/length(zStar))
  }
  else{
    term2<-0
  }
  
  pTerms<-list(term1,term2,term3)
  #pTerms<-cbind(term1,term2,term3)
  
  
  sc<-c(term1,
        term2,
        term3)
  
  label=c(rep("selection", length(term1)),
          rep("immigration", length(term2)),
          rep("transmission", length(term3)))
  
  
  pTerms<-data.frame(species=names(sc),contrib=sc, mechanism=label)
  
  return(pTerms)
}

#Figure 3 computes the KL divergence of several species.


load( file="exampleCode/TannerTable84.RData")
load( file="exampleCode/TannerTable94.RData")


ppSummary<-propPieceWImmigrants(past=data1984,pres=data1994)

ppSCopy<-ppSummary #This copy is used lower down in case of an accidental overwrite of code


deltaPi<-ppSummary$pres$pidecendants-ppSummary$past$pi    


#####Generate Figure 3 illustraing distributions of 3 species##################


pjGiveniPlot<-ppSummary$past$pjGiveni[,colnames(data1984) %in% c("Lyonia octandra","Cyrilla racemiflora","Cinnamomum montanum")]
pjGiveniPlot<-melt(pjGiveniPlot)
pjGiveniPlot<-cbind(pjGiveniPlot,pj=ppSummary$past$pj )
pjGiveniPlot<-melt(pjGiveniPlot)

pjGiveniPlot$Var2= factor(pjGiveniPlot$Var2, levels=c("Cyrilla racemiflora","Lyonia octandra","Cinnamomum montanum"))
pjGiveniPlot$Var1= factor(pjGiveniPlot$Var1, levels=c("col", "w slope","Mull","Mor"))

#rename levels to improve plot readability
levels(pjGiveniPlot$Var1)= c("Col", "Slope","Mull","Mor")
levels(pjGiveniPlot$Var2)= c("A) C. racemiflora","B) L. octandra","C) C. montanum")
levels(pjGiveniPlot$variable)<-c(expression(P[M]),"A")


ggplot(pjGiveniPlot, aes(x=Var1, y=value, fill=variable)) + 
  facet_wrap(~Var2)+
  geom_col(aes(y=value), position="dodge")+ 
  scale_fill_manual(values=c("#7030a0ff", "#0088aaff"), name="Distribution", labels=c(expression(P["j|i"]),expression(P[.j])))+
  xlab("")+
  ylab("Probability")+
  theme(panel.grid.major = element_blank(), panel.grid.minor = element_blank(),
        panel.background = element_blank(), axis.line = element_line(colour = "black"), legend.text = element_text(size = 12))


######Figure 4 bootstrap estimates of all diversity components from 1984-1994#########################


#AS of now this code implements elena's first test to obtain confidence intervals for the significance
#of diversity change and the significance of each component od viersity changej


allTimeBS<-data.frame(condition=factor(),mechanism=factor(),value=numeric(),ci.lo=numeric(),ci.hi=numeric())

#deltaH<-data.frame(gamma=NA,alpha=NA, beta=NA)

  
  d84<-data1984
  d94<-data1994

allTimeBS=rbind(allTimeBS,bootStrapFun(data1984,data1994))
  


#allTimeBS$sampleInterval<-gl(4,12,labels=sampleIntervals)

condition=factor(c("gamma","gamma","gamma","gamma","alpha","alpha","alpha","alpha","beta","beta","beta","beta"), levels=c("gamma","alpha","beta"))


levels(allTimeBS$mechanism)[2]<-"recruitment"
allTimeBS$conditionN= factor(allTimeBS$condition, levels=c('beta','gamma','alpha'))
levels(allTimeBS$conditionN)<-c("A) Beta diversity","B) Gamma diversity","C) Alpha diversity")


ggplot(allTimeBS, aes(x=mechanism, y=value, group=mechanism, color=mechanism)) + 
  facet_wrap(~conditionN)+
  geom_hline(yintercept =  0, size=1)+
  geom_pointrange(aes(ymin=ci.lo, ymax=ci.hi), size=1)+
  theme(panel.grid.major = element_blank(), panel.grid.minor = element_blank(),axis.text.x=element_blank(),
        panel.background = element_blank(), axis.line = element_line(colour = "black"))+
  xlab("")+
  ylab("contribution to diversity change")


###############Monte carlo test for the effects of drift##################


allMC<-data.frame(ci.lo=numeric(),ci.hi=numeric(), observed=numeric(), pvalue=numeric())


  
past<-data1984
present<-data1994
  


source("bootstrap drift test oct 19.R")
  
allMC=rbind(allMC,driftTest)
  
allMC$diversity<-rep(c("gamma","alpha","beta"),1)


#########Summary statistics described in text##############


#in the pre hurricane survey SW beta diversity was 0.53
log(d(data1984,lev="beta",wts=ppSummary$past$pj))

#Beta diversity was essentially unchanged between 1984 and 1994
log(d(data1994,lev="beta",wts=ppSummary$pres$pj))-log(d(data1984,lev="beta",wts=ppSummary$past$pj))


#Gamma diversity incrased substantially
log(d(data1994,lev="gamma",wts=ppSummary$pres$pj))-log(d(data1984,lev="gamma",wts=ppSummary$past$pj))


##########find contributions of each species##########

#We mention that pittosporum undulatum's contribution to gamma diveristy change was far larger
#than the contributions of other immigrant species.

#In the main text, equation 7 sums up the contribution of each species to each mechanism.
#Here we make the mathematicallyu equivalent calcualtion but assesing each species individually.

speciesContrib<-classicPriceBySpecies(ppSCopy$past$pi,       #frequency of ancestors
                                      ppSCopy$pres$pidecendants,      #frequency individuals decendend from ancestral community
                                      ppSCopy$pres$freqofImmigrantpool,      #frequency of each unconnected type
                                      ppSCopy$past$Zgamma,         #traits of ancestors
                                      ppSCopy$pres$Zgamma,         #traits of decendants community
                                      ppSCopy$pres$ZgammaImm,       #trait value of unconnected populations
                                      ppSCopy$pres$pInImmigrantpool) 


#here is the contribution of each immigrant species, including pittosporum undulatum. 
speciesContrib[speciesContrib$mechanism=="immigration",]


#there is fair bit of book keeping the computation above 
#the following code is is a check that it is correct


#here is a computation of the mechanisms analogus to equaiton 7 in the main text
mechanismContrib<-classicPriceExtended1(ppSCopy$past$pi,       #frequency of ancestors
                                      ppSCopy$pres$pidecendants,      #frequency individuals decendend from ancestral community
                                      ppSCopy$pres$freqofImmigrantpool,      #frequency of each unconnected type
                                      ppSCopy$past$Zgamma,         #traits of ancestors
                                      ppSCopy$pres$Zgamma,         #traits of decendants community
                                      ppSCopy$pres$ZgammaImm,       #trait value of unconnected populations
                                      ppSCopy$pres$pInImmigrantpool) 


#Here we sum up each species' contribution
sumOfSpContrib<-aggregate(contrib~mechanism,speciesContrib, sum)

#we get the exact same values (though poorly labeled) when we look at mechanism contributions
mechanismContrib
```
